# CiBER-seq dissects genetic networks by quantitative CRISPRi profiling of expression phenotypes

**DOI:** 10.1101/2020.03.29.015057

**Authors:** Ryan Muller, Zuriah A. Meacham, Lucas Ferguson, Nicholas T. Ingolia

**Affiliations:** Department of Molecular and Cell Biology, University of California, Berkeley, Berkeley, CA 94720, USA; California Institute for Quantitative Biosciences, University of California, Berkeley, Berkeley, CA 94720, USA

## Abstract

To realize the promise of CRISPR/Cas9-based genetics, approaches are needed to quantify a specific, molecular phenotype across genome-wide libraries of genetic perturbations. We address this challenge by profiling transcriptional, translational, and post-translational reporters using CRISPR interference with barcoded expression reporter sequencing (CiBER-seq). Our barcoding approach connects an entire library of guides to their individual phenotypic consequences using pooled sequencing. We show that CiBER-seq profiling fully recapitulates the integrated stress response (ISR) pathway in yeast. Genetic perturbations causing uncharged tRNA accumulation activated ISR reporter transcription. Surprisingly, tRNA insufficiency also activated the reporter, independent of the Gcn2 kinase that senses uncharged tRNAs. By uncovering alternate triggers for ISR activation, we illustrate how precise, comprehensive CiBER-seq profiling provides a powerful and broadly applicable tool for dissecting genetic networks.

## INTRODUCTION

CRISPR-Cas9 has emerged as a powerful and versatile tool for creating precise, programmable genetic perturbations *(1)*. CRISPR-based knockout *(2–4)* and knockdown *(5)* approaches now enable systematic, genome-wide genetic analysis in a wide range of cells and organisms. The Cas9 protein binds short RNAs that guide this protein-RNA complex to complementary sites in the genome, where it can induce mutations or silence promoters *(1)*. Libraries of guide RNAs, each targeting one individual gene, can be used to create a population of cells that each express one distinct guide *(6–8)*. The phenotype of each cell then reflects the impact of the single guide that it expresses. It is straightforward to identify guides that affect cell survival or proliferation, but growth is a crude phenotype that is poorly suited to address many important biological questions *(9)*. The scope of CRISPR-based genetics would be expanded by improved techniques to measure more specific and relevant phenotypes across this diverse population and link these measurements back to individual guides.

Molecular phenotypes, such as the expression level of a critical gene or the stability of a key protein, provide a focused and sensitive gauge for many aspects of cell physiology. We present an approach for profiling a transcriptional, translational, or post-translational regulatory response comprehensively across CRISPR-based perturbations genome-wide. We adapt barcoded expression reporters *(10)* to produce quantitative phenotypic profiles from bulk sequencing of highly diverse populations. These profiles also enable high-precision genetic interaction analyses, which use double mutant phenotypes to map the structure of regulatory networks *(11)*. This direct sequencing approach offers significant advantages over fluorescent reporters for CRISPR-based genetics. Fluorescence phenotypes are typically analyzed by cell sorting *(9)*, which imposes bottlenecks on the cell population size and discretizes quantitative fluorescence measurements into a few broad gates. Our approach circumvents both of these limitations. It also complements broader expression profiles from single-cell approaches such as Perturb-Seq *(12, 13)*, CRISP-Seq *(14)*, and CROP-Seq *(15)*, which cannot capture enough cells to approach genome-scale coverage. These technically challenging studies each analyzed fewer than a hundred genes, and in fact, some relied on fluorescent reporter screens to choose a small, tractable set of guides for single-cell experiments *(12)*. Better ways to profile molecular phenotypes across genome-scale guide libraries thus stand to benefit many areas of biology.

Here, we combine CRISPR interference with barcoded expression reporter sequencing (CiBER-seq) to measure cellular responses provoked by guide RNA-mediated knockdown. Using CiBER-seq, we analyze the integrated stress response (ISR) in budding yeast. We confirm the known pathway for ISR activation by amino acid starvation, which leads to the accumulation of uncharged tRNAs and inhibitory phosphorylation of the translation initiation factor eIF2α *(16)*. Surprisingly, we also find that tRNA depletion triggers the ISR through a distinct, uncharacterized mechanism that does not require eIF2α phosphorylation. We demonstrate an indirect CiBER-seq approach to measure translational and post-translational regulation, and use this approach to show that ISR activation by tRNA depletion nonetheless results from translational control independent of eIF2α phosphorylation. By analyzing double-knockdown phenotypes in dual-guide CiBER-seq, we uncover epistatic interactions that suggest a role for actin cytoskeletal components in mediating this non-canonical ISR. Our results illustrate the power of CiBER-Seq to provide new insights even in well-studied regulatory pathways and highlight the flexibility and broad applicability of our technique.

## RESULTS

### Barcoded expression reporters link transcriptional responses with guide RNA-mediated perturbations in massively parallel screens

CiBER-Seq relies on massively parallel measurements of reporter expression in a diverse population by deep sequencing of short sequence “barcodes” embedded in the reporter transcript *(10)*. The underlying abundance of cells with each barcode can vary in this population, however, and so we set out to measure the transcriptional output from each barcode by normalizing the RNA abundance of that barcode against its DNA abundance. We first validated the use of barcode sequencing to measure expression using an artificial inducible promoter, *P(Z) (17)*, to drive the expression of a barcoded fluorescent reporter, allowing us to vary transcription of the reporter (fig. S1A) and compare fluorescence levels with barcode sequencing results. We generated three high-diversity barcoded libraries, each incorporating a distinct library sequence tag alongside random nucleotide barcodes, and treated these tagged populations with different levels of inducer (Fig. 1A). We measured fluorescence levels in each of these populations (Fig. 1B), and pooled them together to test our ability to measure expression differences in complex populations. The abundance of individual barcodes varied widely in each population (fig. S1D and S1E), but the RNA-to-DNA ratio showed a much tighter distribution that matched quantitatively the fluorescence levels (Fig. 1C).

**Fig. 1.**
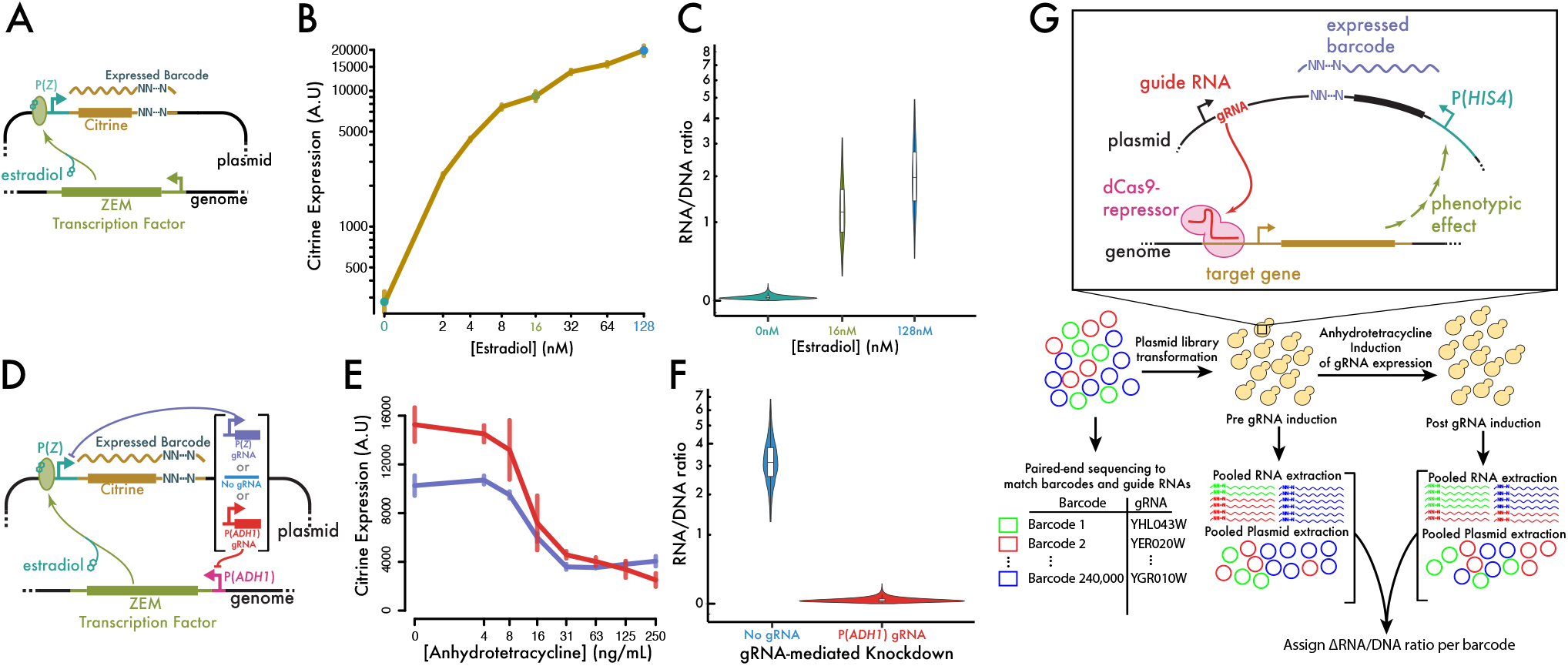
Barcoded expression reporters accurately link guide RNAs with transcriptional phenotypes. (**A**) Schematic of synthetic ZEM transcription factor and *P(Z)* promoter. (**B**, **C**) Expression measurements by (B) fluorescence and (C) RNA/DNA barcode count ratio show quantitative agreement. Inducer conditions for barcode sequencing in (C) are color-coded and marked with dots on the graph in (B). (**D**) Schematic of guide RNA effects on the synthetic transcription factor / promoter pair. (**E**) Fluorescent output is negatively correlated with guide RNA induction level. (**F**) Barcode sequencing expression measurements reflect effects of knockdown. (**G**) Schematic of CiBER-Seq profiling experiment.

Having established our ability to track transcriptional output through barcode sequencing, we next sought to link distinct barcoded reporters with guide RNAs that perturb reporter expression (Fig. 1D). The inducible promoter driving our reporter relies on an exogenous, chimeric transcription factor, ZEM (17). CRISPRi-mediated knockdown of this transcription factor, or the inducible P(Z) promoter itself, both reduced reporter fluorescence dramatically (Fig. 1E and fig. S1B and C). We then carried out pooled sequencing on two high-complexity barcode libraries, one linked to the guide RNA that targeted the exogenous transcription factor. Barcode sequencing revealed a 91-fold reduction in reporter expression caused by transcription factor knockdown (Fig. 1F), demonstrating our ability to measure CRISPRi effects through a linked, barcoded reporter.

Based on these results, we devised a strategy for genome-wide CRISPR interference with barcode expression reporter sequencing (CiBER-seq). We linked each guide sequence in a comprehensive yeast guide RNA library with random nucleotide barcodes and determined the linkage between barcodes and guides by paired-end sequencing (Fig. 1G) *(18)*. Our library contains 10 guides per gene (~60,000 in total) and ~240,000 distinct barcodes (~4 per guide on average). By linking multiple barcodes with each guide, we were able to obtain independent measurements of guide effects within a single experiment. To control for barcode-specific effects on RNA stability, we drove guide RNA expression from a tetracycline-inducible promoter *(19)* and compared barcode expression before and after guide induction. Inducible CRISPRi also facilitates measurements of guides with strong fitness effects by allowing us to propagate cells without guide induction and thereby avoid the premature loss of guides with growth defects. Finally, we noted that barcode DNA-seq showed substantially more noise than barcode RNA-seq (fig. S1D and S1E), which we attribute to PCR amplification. In order to reduce this noise and improve the precision of our expression measurements, we constructed subsequent DNA libraries using linear pre-amplification by in vitro transcription *(18)*.

### CiBER-seq recapitulates known genetic regulators of integrated stress response and identifies new regulators related to tRNA insufficiency

We first applied CiBER-Seq to survey genetic perturbations that activated a transcriptional reporter for the yeast integrated stress response (ISR), also known as the general amino acid control (GAAC) response. This deeply conserved pathway upregulates biosynthetic genes in response to elevated levels of uncharged tRNAs that arise during amino acid starvation *(16)*. We constructed a guide library with barcodes driven by the promoter *P(HIS4)*, which is activated by the ISR transcription factor Gcn4 when uncharged tRNAs accumulate. In order to distinguish ISR-mediated regulation from nonspecific or global effects on transcription, we also constructed a second guide library using the housekeeping promoter *P(PGK1)*. CiBER-Seq profiling of *P(HIS4)* induction identified 35 different guide RNAs targeting amino acid biosynthesis pathways, along with guides against 17 of the 20 of the aminoacyl-tRNA synthetases (Fig. 2, A and C), which all interfere with tRNA charging. We also observed *P(HIS4)* activation in response to knockdown of each individual component of the eIF2 translation initiation complex (Fig. 2, A and C); depletion of these proteins directly increases translation of the Gcn4 transcription factor, leading to *P(HIS4)* induction *(16, 20)*. All of these CRISPRi perturbations affected *P(HIS4)* specifically, and did not induce *P(PGK1)* (Fig. 2B), although knockdown of general RNA polymerase II transcription reduced expression in both reporters (fig. S2, A through C). In fact, *P(PGK1)*-driven barcode expression was induced instead by knockdown of other glycolytic enzymes (fig. S2, A through C), suggesting a homeostatic transcriptional activation of *PGK1* expression in response to impaired glycolysis.

**Fig. 2.**
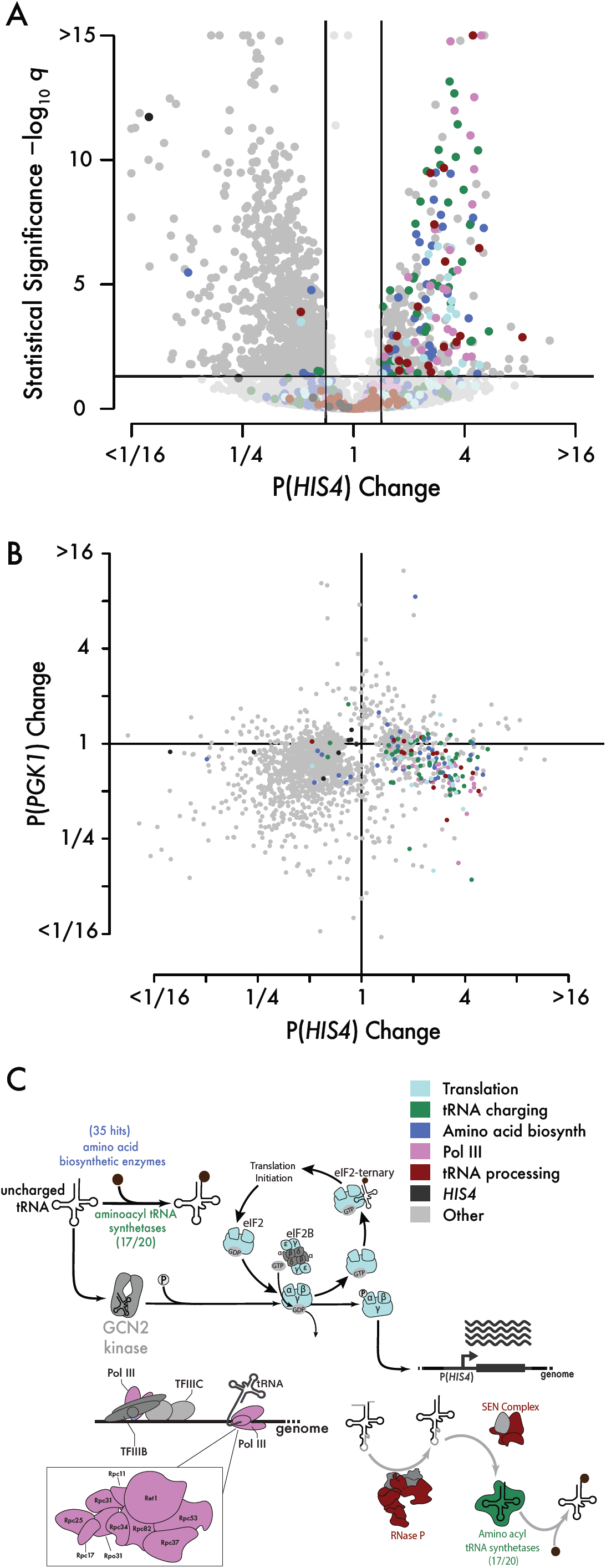
CiBER-seq recapitulates known genetic regulators of integrated stress response and identifies new regulators related to tRNA insufficiency. (**A**) Genome-wide CiBER-seq profile of P(*HIS4*) transcription. Each point represents a different guide RNA, colored according to the function of the target gene. The lines indicate guides with a significant (*q* < 0.05) and substantial (>1.5-fold change) effect. (**B**) Comparison of CiBER-seq profiles of P(*HIS4*) and *P(PGK1)* transcription, colored as in (A). (**C**) Coverage of the known integrated stress response pathway, RNA polymerase III, and tRNA processing complexes by guides with significant effects on *P(HIS4)* activity.

Surprisingly, we also found that overall tRNA depletion triggered the ISR transcriptional program. Guides targeting subunits of RNA polymerase III, which transcribes tRNAs, as well as the tRNA processing complex RNAse P and the SEN tRNA splicing complex *(21)* all activated *P(HIS4)* transcription (Fig. 2, A and C). In contrast to known ISR triggers, these genetic perturbations should not lead to the accumulation of uncharged tRNAs and in fact should reduce overall tRNA levels. While depletion of initiator methionyl-tRNA can induce *P(HIS4) (22)* and RNA polymerase III defects can lead to initiator methionyl-tRNA depletion *(23)*, this effect cannot explain our observations here. Initiator tRNA does not contain an intron, and so the loss of SEN should not reduce initiator tRNA levels. To further exclude initiator tRNA depletion as an explanation for the effects of RNA polymerase III knockdown, we overexpressed initiator tRNA during CRISPRi-mediated ISR activation (fig. S2J and S2K). Impaired tRNA transcription showed no particular susceptibility to suppression of ISR response by high-copy overexpression of initiator tRNA, relative to disruption of amino acid biosynthesis or tRNA charging (fig. S2K). Taken together, our results argue that elongator tRNA depletion can directly activate ISR transcription, perhaps through its effects on translation elongation.

### tRNA insufficiency triggers HIS4 transcription independently of eIF2a phosphorylation and Gcn2 kinase

Because the integrated stress response is not activated during log-phase growth, our initial CiBER-seq data did not identify expression changes caused by loss of canonical, positive transducers of ISR signaling, such as *GCN2* and *GCN4*. To uncover the phenotypes of these and other ISR pathway genes, we next looked for regulators whose depletion would block *P(HIS4)* activation triggered by the toxic histidine analog 3-amino-1,2,4-triazole (3AT), which blocks histidine biosynthesis (Fig. 3, A and B). Guides that activate *P(HIS4)* under replete nutrient conditions did not further elevate *P(HIS4)* expression upon 3AT treatment (Fig. 3B). We infer that Gcn4-mediated activation is saturated under these conditions, although we note that knockdown of Gcn4 degradation factors, such as *PCL5 (24)*, can enhance 3AT-mediated *P(HIS4)* induction. Meanwhile, guides that reduced *P(HIS4)* transcription, including guides targeting the core RNA polymerase II transcription machinery (fig. S2, A, B, and C), did not interfere with *P(HIS4)* activation upon 3AT treatment. We therefore reasoned that guides affecting amino acid biosynthesis, as well as tRNA charging, transcription, and processing, each saturated *P(HIS4)* transcription through a shared, *GCN4*-dependent mechanism. More broadly, these results illustrate the power of CiBER-Seq to dissect gene regulatory networks and identify novel interactors in even well-characterized systems.

**Fig. 3.**
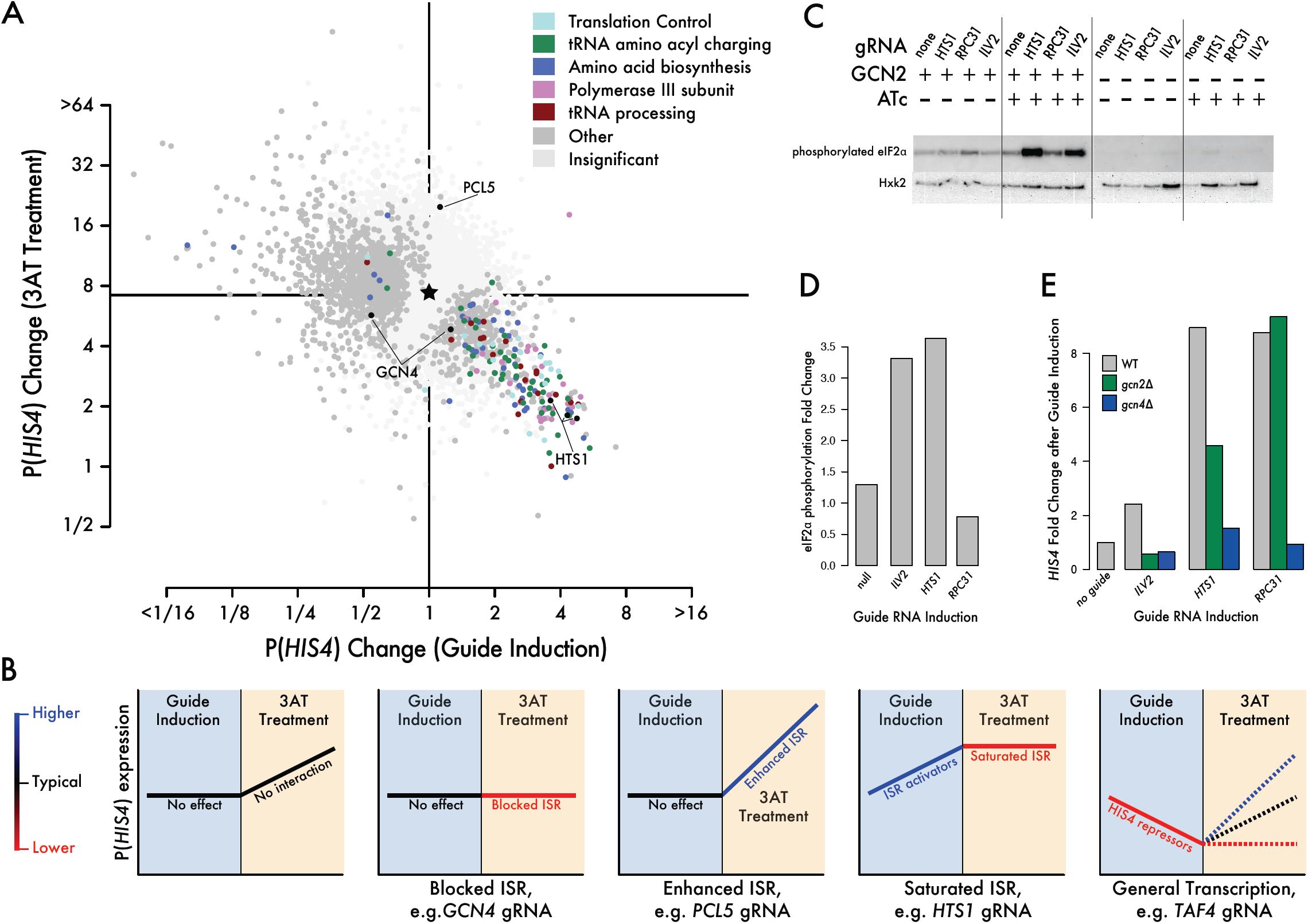
tRNA insufficiency activates HIS4 transcription through a pathway that bypasses GCN2 kinase and eIF2a phosphorylation. (**A**) Comparison of P(HIS4) CiBER-seq profiles before and after 3AT treatment. (**B**) Schematic outlining observed ISR responses for functional categories of genes. (**C**) Western blot for eIF2α phosphorylation. Knockdown of amino acid biosynthesis (*ILV2* guide) and aminoacyl tRNA charging (*HTS1* guide) results in eIF2α phosphorylation while knockdown of RNA polymerase III (*RPC31* guide) does not. *GCN2* deletion abolishes eIF2α phosphorylation. (**D**) Quantification of eIF2α phosphorylation relative to hexokinase loading control in (C). (**E**) Endogenous *HIS4* mRNA levels measured by qPCR. Endogenous *HIS4* activation by *ILV2, HTS1,* or *RPC31* knockdown is completely *GCN4* dependent. *HIS4* activation by *ILV2* knockdown is also completely *GCN2* dependent while effects of *RPC31* knockdown are entirely *GCN2* independent and *HTS1* knockdown is only partially *GCN2* dependent.

Since one hallmark of ISR activation is the phosphorylation of eIF2α, we were curious whether tRNA insufficiency provoked this response. While CRISPRi knockdown of the amino acid biosynthetic enzyme *ILV2* or the histidyl-tRNA synthetase *HTS1* both induced eIF2α phosphorylation, knock-down of the RNA polymerase III subunit *RPC31* did not (Fig. 3C and 3D), consistent with previous observations *(23)*. Deletion of *GCN2*, the only known eIF2α kinase in yeast, completely blocked eIF2α phosphorylation in all conditions tested (Fig. 3C). Activation of *P(HIS4)* in the absence of eIF2α phosphorylation further distinguished the response to tRNA depletion upon *RPC31* knockdown and the classical ISR pathway.

We thus tested the genetic requirements for activation of endogenous *HIS4* transcription in response to these CRISPRi-mediated perturbations. Deletion of *GCN4* blocked *HIS4* activation by *RPC31* knockdown as well as *ILV2* and *HTS1* knockdown, and so the effects of tRNA insufficiency reflect a *GCN4-*mediated ISR. Strikingly, *GCN2* deletion produced distinct effects across these three CRISPRi knockdowns (Fig. 3E). Deletion of *GCN2* completely blocks *HIS4* induction in response to *ILV2* depletion, consistent with canonical models of ISR signaling (25). Although *HTS1* knockdown triggers strong eIF2α phosphorylation, *GCN2* deletion only partially abrogated its transcriptional effects. Finally, *RPC31* knockdown induces *HIS4* transcription independent of *GCN2*, as we expected based on the lack of eIF2α phosphorylation. These results suggest that translation elongation defects, arising from impaired tRNA recruitment, directly trigger *GCN4*-mediated transcriptional responses. Knockdown of tRNA synthetases activates the ISR by this pathway in parallel with the *GCN2*-dependent response to uncharged tRNAs.

### Perturbations of the ARP2/3 complex prevent ISR activation by HTS1 or RPC31 knockdown

Given that both Gcn2 activity and eIF2α phosphorylation were dispensable for ISR activation in response to tRNA depletion, we next sought to systematically identify genes required for this response. We looked for genetic perturbations that modified *P(HIS4)* responses to different ISR triggers by performing dual-guide CiBER-Seq, combining one guide that individually activated the ISR with a second guide from our genome-wide CRISPRi library to obtain quantitative, genome-wide genetic interaction profiles. We compared the interaction profiles of the histidyl-tRNA synthetase *HTS1*, which is required for tRNA charging, with those of the RNA polymerase III subunit *RPC31*, which is required for tRNA transcription (Fig. 4, A, B, and C). We saw saturated *P(HIS4)* induction after knockdown of *HTS1* or *RPC31,* similar to the *P(HIS4)* saturation we observed upon 3AT treatment (Fig. 3A and 3B), as guides that activated the ISR on their own did not further enhance transcription in these dual-guide epistasis experiments. We also observed guides that had no effect on their own, but either suppressed ISR activation, such as guides against *GCN4* itself, or enhanced the strength of the response, including guides against the degradation factor *PCL5* or the ISR inhibitor *YIH1 (26, 27)*.

**Fig. 4.**
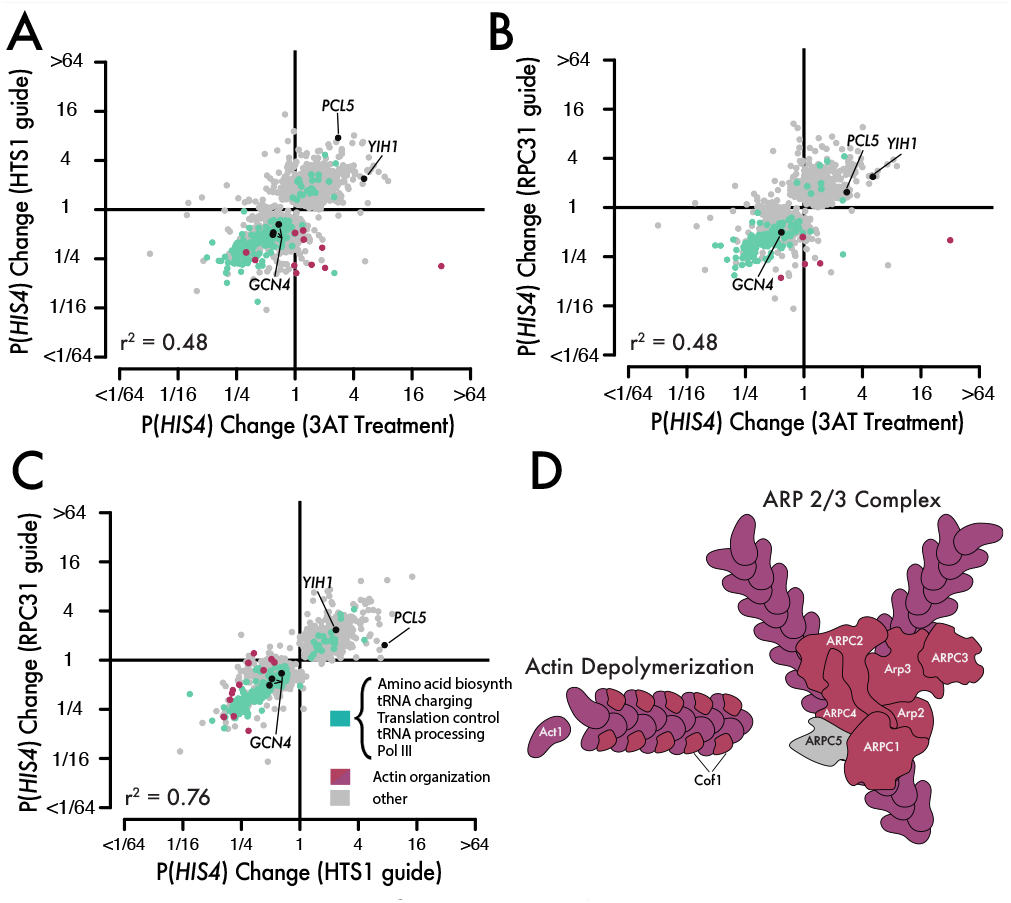
Perturbations of the ARP2/3 complex prevent ISR activation by HTS1 or RCP31 knockdown. (**A** to **C**) Pairwise comparison of *P(HIS4)* CiBER-seq profiles between ISR activation by 3AT treatment, *HTS1* knockdown, and *RPC31* knockdown. Each CiBER-seq profile was generated by measuring changes in *P(HIS4)* expression between the the guide in the context of an ISR activator relative to the change cause by the guide in isolation (Fig. 2A). (**D**) Coverage of actin cytoskeleton and ARP2/3 complex by guides that block ISR induction during *HTS1* or *RPC31* knockdown, but not 3AT treatment.

We compared the genetic interaction profiles across three different ISR stimuli to test whether they were acting through similar or distinct pathways. Epistatic profiles of *HTS1* and *RPC31* knockdown resembled each other (*R^2^* = 0.76) more closely than either resembled the profile during 3AT treatment (*R^2^* = 0.48), although all three did overlap substantially. This pattern aligns with our observation that *P(HIS4)* activation by either tRNA charging defects or tRNA depletion are not completely *GCN2* dependent, whereas the amino acid starvation response is fully *GCN2* dependent (Fig. 3E). We thus looked for genetic interactions specific to these *GCN2*-independent responses and identified a surprising requirement for the actin cytoskeletal components in response to *HTS1* and *RPC31* knockdown (Fig. 4). Notably, knockdown of actin itself (*ACT1*) or genes encoding members of the Arp2/3 complex did not affect P(*HIS4*) transcription in normal growth or 3AT treatment (fig. S4A), but blocked P(*HIS4*) activation by tRNA charging defects or tRNA depletion (fig. S4, B and C). This could reflect interactions between the ISR inhibitor *YIH1* and free actin monomers (27), or instead arise because of nuclear actin’s role in transcription *(28)*.

### Sumoylation is a post-translational regulator of Gcn4

CiBER-Seq analysis of *P(HIS4)* regulation provided a comprehensive view of yeast ISR signaling, revealing distinct triggers for *GCN4*-dependent transcriptional responses. However, this approach does not allow us to distinguish the different layers of Gcn4 regulation. Gcn4 abundance is controlled by regulated protein degradation *(29)* in addition to its well-characterized translational induction upon eIF2α phosphorylation *(16)*. Indeed, we saw epistatic enhancement of ISR activation from guides depleting the decay factor *PCL5*. We thus wanted to specifically analyze the regulators of Gcn4 stability and translation, in isolation, and separate them from other effects on *P(HIS4)* activity. In order to couple these protein-level phenotypes with a transcriptional readout suitable for CiBER-seq, we returned to the synthetic, ZEM chimeric transcription factor used to initially validate barcode sequencing. Since barcode transcription is directly linked to the abundance of this synthetic transcription factor (Fig. 1B), we reasoned that it could couple protein-level regulation with expressed RNA barcode abundance (Fig. 5A).

**Fig. 5.**
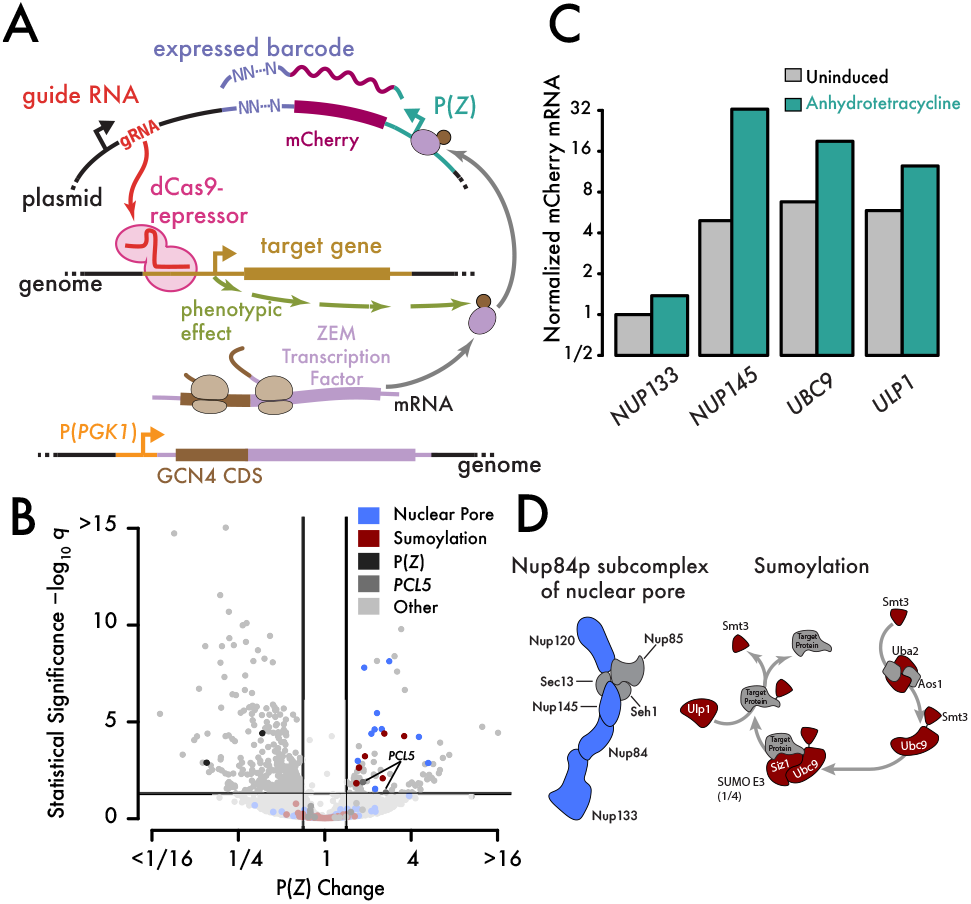
Post-translational regulation of Gcn4 by sumoylation. (**A**) Schematic of indirect CiBER-seq experiment to identify post-translational regulators of Gcn4 (**B**) CiBER-seq response of Gcn4-ZEM / *P(Z)* as in Fig. 2A. (**C**) Measurement of *mCherry* reporter mRNA abundance by qPCR. Knockdown of *NUP133, NUP145, UBC9*, or *ULP1* increase reporter mRNA levels.(**D**) Guides that activate *P(Z)* with Gcn4-ZEM map to the Nup84 nuclear pore subcomplex and to the sumoylation pathway.

We first profiled post-translational control of Gcn4 by carrying out CiBER-Seq in yeast that constitutively express a protein fusion between Gcn4 and the chimeric ZEM transcription factor (Fig. 5A). Levels of this transcription factor should reflect only post-translational regulation of Gcn4, and not its translational regulation, as it is expressed using the *PGK1* promoter and 5′ UTR. Knockdown of *PCL5*, which regulates Gcn4 degradation, affected the activity of the Gcn4-ZEM post-translational reporter. Neither tRNA charging nor tRNA biogenesis had an effect, however. Instead, we found that disruption of sumoylation increased expression from *P(Z*) (Fig. 5, B and C). We identified individually significant guide RNAs targeting nearly every step of the sumoylation pathway, including SUMO itself, SUMO-activating and SUMO-conjugating enzymes, and the Siz1 SUMO ligase *(30)*. Likewise, we identified guides against most components of the Nup84 subcomplex of the nuclear pore *(31)* that increased Gcn4-ZEM activity. This broad coverage of protein complex subunits and functionally related genes, here in the SUMOylation pathway and the Nup84 subcomplex (Fig. 5C), highlights the value of comprehensive, quantitative CiBER-Seq profiling. Sumoylation of Gcn4 has been shown to promote its eviction from chromatin, even when recruited through heterologous DNA-binding domains, as well as its subsequent degradation, both of which are expected to limit Gcn4-ZEM transactivation of barcoded reporters *(32, 33)*. Knockdown of the SUMO conjugation machinery would thus relieve these limiting effects and enhance barcode expression. Elevated barcode expression during knockdown of the

Ulp1 deconjugating protease could result from its role in SUMO maturation *(34)* or reflect a more complex requirement for a sumoylation-desumoylation cycle in Gcn4 regulation. The role of the nuclear pore is likely linked to sumoylation. In yeast, Ulp1 binds physically to the nuclear pore *(35, 36)*, and mutations in the Nup84 complex can mimic some *ulp1* phenotypes through mislocalization *(37)* and reduction of Ulp1 activity *(38)*. Furthermore, Ulp1 at the nuclear pore has been implicated directly in transcriptional activation *(39)*.

### The GCN4 5′ leader sequence is an intrinsic biosensor of translation stress

We observed clear and coherent patterns of genetic perturbation affecting Gcn4 protein activity, but no evidence that tRNA insufficiency affected its post-translational regulation. We next explored the translational control of *GCN4*, which results from regulatory upstream open reading frames in the *GCN4* 5′ leader sequence *(16)*. In order to capture perturbations that affect this translation regulation, we drove expression of the ZEM synthetic transcription factor from the *GCN4* promoter and 5′ leader sequence and assessed guide effects on barcode expression (Fig. 6A).

**Fig. 6.**
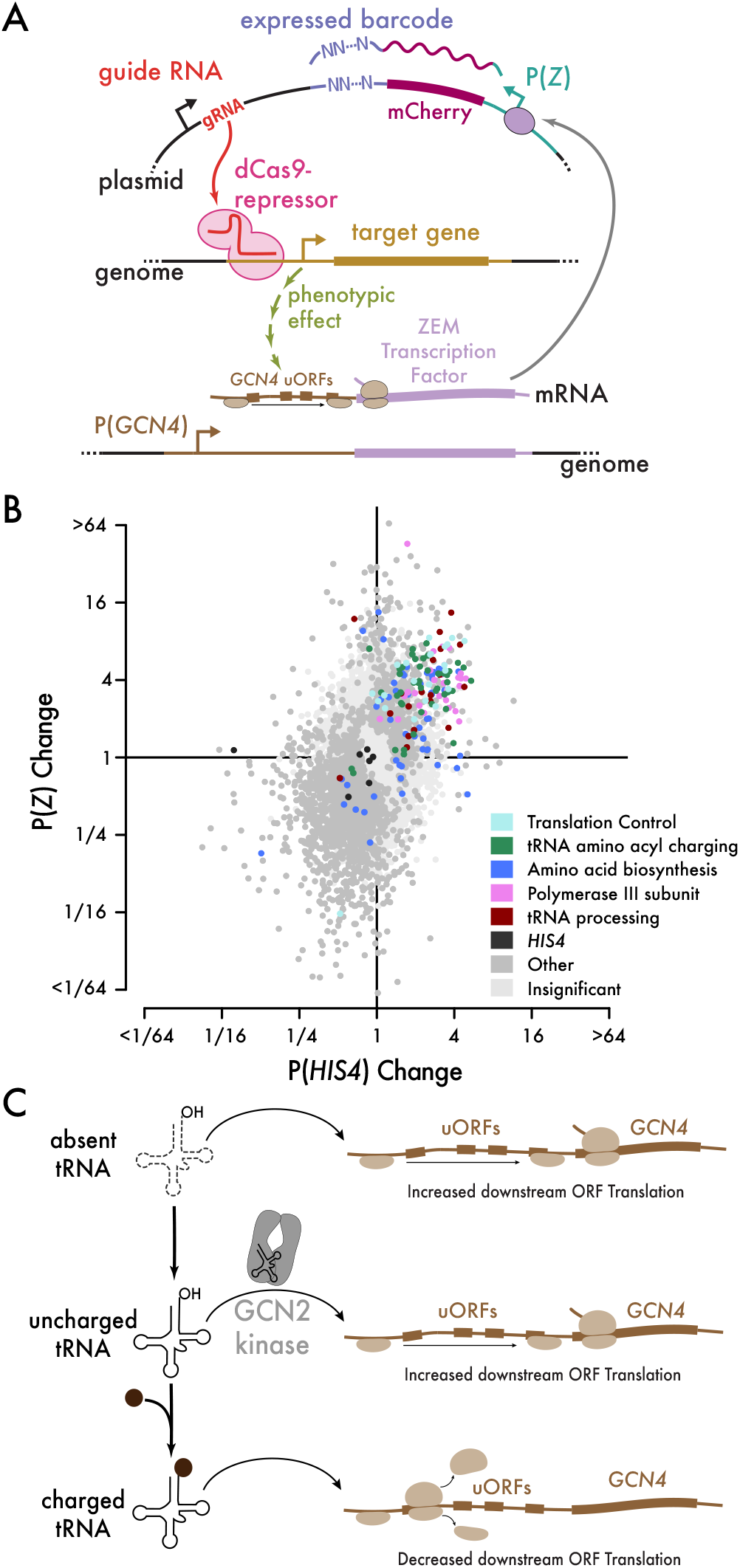
The GCN4 5′ leader sequence is an intrinsic biosensor of translation stress. (**A**) Schematic of indirect CiBER-seq experiment to identify mediators of *GCN4* 5′ leader translation regulation. (**B**) Comparison of direct *P(HIS4)* CiBER-seq profile versus indirect *P(Z*) profile, with ISR activators colored according to their function. (**C**) Model of *GCN4* 5′ leader sequence as an intrinsic biosensor of translation stress.

The same guides that activated *P(HIS4)*, including those that block tRNA biogenesis as well as tRNA charging, also increased *P(Z)* in our CiBER-seq analysis of translational control through the *GCN4* 5′ leader (Fig. 6B). By assessing this translational response in isolation, we observe effects occur in the same direction, but with larger magnitude, than *P(HIS4)*. This analysis supports the view that translational regulation is central to ISR activation, even when it proceeds independently of *GCN2* and eIF2α phosphorylation, although our data cannot exclude transcriptional effects mediated by the *GCN4* promoter *(40)*.

## DISCUSSION

By linking CRISPR interference guides with barcoded expression reporters, we generate quantitative, genome-wide phenotypic profiles for specific molecular events in the cell. Our CiBER-seq approach allows us to address transcriptional, translational, and post-translational regulation, enabling systematic genetic analysis of diverse biological processes. We leveraged quantitative CiBER-seq phenotypic profiles to gain new and surprising insights into the integrated stress response, identifying tRNA depletion as a previously unappreciated trigger for this well-characterized pathway.

Because translation is a resource-intensive biosynthetic process, most organisms sense translational stresses and respond with physiological changes that maintain homeostasis. In eukaryotes, amino acid starvation, which directly impacts tRNA charging and translation elongation, triggers a global decrease in translation initiation while also increasing transcription of amino acid biosynthesis genes *(16)*. In many cases, reduced initiation is sufficient to restore normal elongation profiles *(41)*. Indeed, when aminoacyl-tRNA synthetases are depleted, yeast reduce translation initiation until tRNA charging and utilization are balanced. Furthermore, in normal circumstances, these synthetases appear to buffer tRNA levels by sequestering uncharged tRNAs *(42)*. Here, we find that the opposite situation — tRNA depletion in the presence of adequate synthetases and amino acids — triggers a *GCN2*-independent ISR. Our comprehensive genetic data implicating many stages of tRNA biogenesis argue that *GCN4* translation responds directly to elongator tRNA insufficiency, rather than depletion of initiator tRNA *(23)* or the accumulation of unprocessed nuclear tRNA precursors *(43)*. As we have not identified other trans-acting regulatory pathways under these circumstances, we propose that the *GCN4* 5′ leader is an intrinsic biosensor for translation elongation stress (Fig. 6C). Artificial stimuli, including tRNA depletion, can activate *GCN4* translation independent of eIF2α phosphorylation. In the natural history of budding yeast, the ISR likely evolved to sense nitrogen starvation and elicit general inhibition of translation through *GCN2*, along with *GCN4*-dependent activation of homeostatic transcriptional programs *(16)*. Our work thus expands the range of stresses known to activate the ISR and points to a general mechanism for sensing translational perturbations.

While we showed that the ISR is activated by artificial, genetic depletion of tRNAs, similar situations may arise in a natural context *(44)*. Starvation and other stresses can induce tRNA cleavage *(45, 46)*, and while much attention has focused on the positive roles of the resulting fragments (47), cleavage can also deplete the tRNA substrate and alter translation *(48)*. Effective tRNA depletion may also arise when tRNAs are sequestered in the nucleus and thus unavailable for cytosolic translation *(21)*. Finally, an array of human disease mutations affecting tRNA biogenesis factors lead to neurodegenerative disorders, although the molecular basis for this effect is not clear *(44, 49)*. Mutations in tRNA genes themselves can lead to tRNA insufficiency, ISR activation, and neurodegeneration, although in mammals this effect is *GCN2* dependent *(50, 51)*.

Our results demonstrate the power of CiBER-seq in elucidating the genetic architecture of regulation in the cell. We see comprehensive coverage of pathways and molecular complexes by guides targeting each individual component, indicating that our approach is both specific and sensitive. We identified ISR activation upon knockdown of each subunit of RNA polymerase III along with most proteins involved in tRNA processing, revealing tRNA depletion as the underlying trigger for the ISR. We identify other modes of regulation as well, such as sumoylation of Gcn4, and even detect the compensatory *P(PGK1)* activation in response to impaired glycolysis. Quantitative CiBER-seq profiles enable genetic interaction analysis that provides further insight into regulatory networks. Correlated patterns of epistasis are a powerful tool for identifying genes that function together in pathways and complexes, and likewise chemogenomic comparison between chemical and genetic interaction profiles can reveal functional drug targets *(11)*. Here, we leverage this advantage of CiBER-seq to identify a role for actin cytoskeletal components in ISR signaling. More broadly, our results highlight the power of CiBER-Seq to combine specific and quantitative molecular phenotypes with targeted genetic perturbations and thereby precisely dissect regulatory pathways. Indeed, a similar ReporterSeq technique was developed simultaneously with this work, and applied to elucidate known and novel pathways linking diverse stressors to the yeast heat shock response *(52)*.

As the key components of CiBER-seq translate into nearly any organism, we anticipate many biological insights arising from broad application of our approach.

## ACKNOWLEDGEMENTS

We thank G. Brar, L. Lareau, E. Ünal, and members of the Ingolia and Lareau labs for discussion and advice. This work was funded by NIH grants DP2 CA195768 and R01 GM130996 (N.T.I.). This work used the Vincent J. Coates Genomics Sequencing Laboratory at UC Berkeley, supported by NIH instrumentation grant S10 OD018174.

## AUTHOR CONTRIBUTIONS

R. M. and N. T. I. designed the experiments. R. M., Z. A. M, and L. F. carried out the experiments. R. M., L. F., and N. T. I. analyzed high-throughput sequencing data. R. M. and N. T. I. conceived the study and wrote the manuscript with input from all authors.

## COMPETING INTERESTS

The authors declare no competing interests.

## DATA AND MATERIALS AVAILABILITY

All high-throughput sequencing data are available from the NCBI SRA under BioProject PRJNA578818. All source code is available at https://github.com/ingolia-lab/CiBER_seq. Plasmids and libraries will be made available through AddGene.

## MATERIALS AND METHODS

### Plasmid materials

pRS416-dCas9-Mxi1 + TetR + pRPR1(TetO)-NotI-gRNA was a gift from Ronald Davis (Addgene plasmid # 73796). pKT0139 was a gift from Kurt Thorn (Addgene plasmid # 8731). pHES836 was a gift from Hana El-Samad (Addgene plasmid # 89195). pHES795 was a gift from Hana El-Samad (Addgene plasmid # 87943). pCfB2337 was a gift from Irina Borodina (Addgene plasmid # 67555). pCfB2226 was a gift from Irina Borodina (Addgene plasmid # 67533). pCfB2189 was a gift from Irina Borodina (Addgene plasmid # 67532).

### Plasmid construction

Plasmid assembly was carried out using standard molecular biology techniques as described below, and verified by Sanger sequencing. All PCR reactions were performed using Q5 polymerase (NEB M0491S) according to manufacturer protocols. Restriction enzymes were obtained from NEB and high-fidelity (HF) variants were used when available. DNA fragments were size-selected by gel electrophoresis in 1% agarose. Gibson assembly reactions were carried out using NEBuilder HiFi DNA Assembly Master Mix (NEB E2621L) with DNA fragments containing homology arms between 15 and 20 base pairs long. DNA was purified and concentrated as necessary with DNA Clean & Concentrator (Zymo D4013). Transformations were performed in Stbl3 chemically competent cells provided by the QB3 Berkeley Macrolab facility and plated on appropriate antibiotic plates for colony selection. Plasmid DNA was purified from liquid cultures using DNA miniprep kit (NEB #T1010).

<u>pNTI660</u> was constructed in several steps. First, pNTI601 (pRS416-dCas9-Mxi1 + TetR + pRPR1(TetO)-NotI-gRNA, AddGene #73796) *(19)* was amplified with NM717 and NM724 *(53)* to isolate the gRNA expression cassette and the resulting PCR product was re-circularized by Gibson assembly to create pNTI646. pNTI646 was then digested with MfeI and combined with oligonucleotide NI1025 to create pNTI660.

<u>pNTI725</u> was created in several steps. First, the promoter *P(PGK1)* was amplified from *S. cerevisiae* BY4741 genomic DNA using NI553 and NI554, and the resulting product was subcloned and used as a template for *P(PGK1)* amplification. Yeast-optimized yECitrine coding sequence was amplified from pNTI189 (pKT0139 AddGene #8731) *(54)* using primers RM151 and RM155 and cloned downstream of *P(PGK1)*. Next, the *P(PGK1)*-yECitrine expression fragment was isolated by digestion with SacI and BsrgI restriction sites, and pNTI660 was linearized by digestion with MfeI. Doublestranded DNA splints were created by annealing oligonucleotides RM317 and RM318, and oligonucleotides RM319 and RM320. Digested *P(PGK1)*-yECitrine and pNTI660 were joined together using these splints in a four-piece Gibson assembly reaction. The NotI site for gRNA insertion was replaced with an NruI site by digesting the backbone with NotI and then introducing single-stranded oligonucleotide RM321 by Gibson assembly. Finally, the Illumina TruSeq Read1 site was added to the beginning of the terminator *T(ADH1*) by annealing singlestranded oligonucleotides RM323 and RM324, extending them using Q5 polymerase, digesting the backbone with AscI and introducing the double-stranded product into the digested plasmid by Gibson assembly.

<u>pNTI726</u> was constructed by replacing *P(PGK1)* in pNTI725 with estradiol-responsive P(*GAL1*). P(*GAL1*) with an embedded Zif268 binding site was PCR amplified from pHES836 using primers RM348 and RM349. pNTI725 was digested with SacI and EcoRI and amplified P(GAL1) was then Gibson assembled into the backbone.

<u>pNTI727</u> was generated by inserting the gRNA sequence targeting *P(Z*) into pNTI726. Plasmid pNTI726 was digested with NruI, singlestranded oligonucleotides RM389 and RM390 were annealed, extended using Q5 DNA polymerase, and the double-stranded product was introduced into the linearized plasmid by Gibson assembly.

<u>pNTI728</u> was generated by inserting the gRNA sequence targeting *P(ADH1*) into pNTI726, using the same methods as pNTI727, using oligonucleotides RM383 and RM384.

<u>pNTI729</u> was constructed by first amplifying the ZEM transcription factor from pNTI638 (pHES795 Addgene #87943) *(17)* in two pieces, using oligonucleotide primers RM352 with RM353 and RM354 with RM355. The easy clone vector pCfB2337 (Addgene #67555) *(55)* was then digested with HindIII and the three fragments were combined by Gibson assembly.

<u>pNTI730</u> was constructed by replacing the promoter *P(ADH1*) in pNTI729 with the *P(PGK1)* promoter. pNTI729 was digested with PvuII and NheI, *P(PGK1)* was amplified from pNTI725 using primers RM417 and RM418, and the two fragments were combined by Gibson assembly.

<u>pNTI731</u> through pNTI737, expressing guide RNAs against *ILV2, HTS1, RPC31, NUP133, NUP145, ULP1*, and *UBC9* respectively, were subcloned into the integrating plasmid pCfB2189. First, DNA fragments containing the guide RNA expression cassette were amplified from the pool of gRNA plasmids using forward primers RM639, RM517, RM518, and RM641 through 644, respectively, that anneal to the unique nucleotide barcode sequence, and a reverse primer RM519 that binds downstream of the gRNA scaffold. Fragments were amplified with Gibson homology arms on each end. The integration vector pCfB2189 (Addgene #67532) *(55)* was digested with HindIII and gRNA-expressing fragments were transferred into this backbone by Gibson assembly.

<u>pNTI738</u> for initiator methionyl-tRNA overexpression was constructed by amplifying the initiator methionyl-tRNA locus *IMT4* from yeast genomic DNA using primers RM636 and RM637, digesting the high-copy yeast plasmid pRS426 (ATCC #77107) *(56)* with EcoRI, and combining them by Gibson assembly. pNTI739 was constructed from pNTI730. The *GCN4* CDS was amplified from yeast genomic DNA using primers RM515 and RM516 and inserted upstream of, and in frame with, the synthetic transcription factor by digesting pNTI730 with NheI and introducing *GCN4* by Gibson assembly.

<u>pNTI740</u> was constructed by digesting NTI730 with PvuII and NheI, amplifying the *GCN4* promoter and 5′ UTR from yeast genomic DNA using primers RM513 and RM514, and combining the two fragments by Gibson assembly.

<u>pNTI741</u>, the *P(HIS4)*-yECitrine PCR template used for downstream CiBER-seq plasmid library preparation, was generated by replacing the *P(PGK1)* in pNTI725 with *P(HIS4)*. pNTI725 was digested with SacI and NheI, *P(HIS4)* was amplified from yeast genomic DNA using primers RM499 and RM500, and the two fragments were combined by Gibson assembly. pNTI742 was made in several steps. First, the NotI restriction site of pNTI660 was replaced with an AvrII restriction site by digesting with NotI and introducing a replacement cassette, comprising annealed oligonucleotides RM459 and RM460, by Gibson assembly. Next, the vector was digested with SacI and MfeI. The *T(ADH1)* terminator and TruSeq R1 priming site were amplified from pNTI726 using primers RM489, RM490, and RM491, in a nested PCR approach where RM490 was used at 0.1x the normal concentration. The resulting PCR product was joined with the digested vector by Gibson assembly.

### Plasmid library construction

All pooled plasmid libraries were constructed by Gibson-style assembly using HiFi DNA assembly master mix (NEB E2621L) and transformed using either ElectroMAX DH10B (ThermoFisher #18290015) or NEB^®^ 10-beta Competent E. coli (NEB #C3019H) according to manufacturer protocol. All column purifications were performed using Zymo DNA Clean and Concentrator-5 (Zymo #D4013). Dilutions of reach transformation were plated to estimate library size and ensure sufficiency library diversity. Plasmid libraries were harvested from liquid culture using the Monarch^®^ Plasmid DNA Miniprep Kit (NEB #T1010). Miniprep kit reagents were scaled to accommodate one spin column for every 5mL of culture.

To generate estradiol-inducible barcode-expressed plasmid libraries (Fig1), pNTI726 and pNTI728 were first digested with BamHI. Five barcodes were generated by annealing primer RM396 with one of five barcode primers, RM391, RM392, RM393, RM394, or RM399, and extending the annealed duplex using Q5 polymerase. Four of these barcodes were inserted by Gibson assembly into BamHI-linearized pNTI726 and the last barcode DNA fragment inserted into BamHI-linearized pNTI728. Gibson reactions were purified and concentrated, transformed into ElectroMAX DH10B cells, and inoculated directly into 200 ml of LB-carbenicillin media. Pooled libraries were harvested when batch liquid culture reached OD of 4.

To generate gRNA plasmid libraries, 100 pg of CRISPRi guide RNA oligonucleotide library was amplified by PCR using Q5 polymerase using primers NM636 and NM637 (53). pNTI742 was digested with AvrII and the amplified guide fragments were introduced into the digested vector by Gibson assembly. The assembly product was purified and transformed into ElectroMAX DH10B cells by electroporation. Electroporated cells were inoculated directly into 500mL of LB-carbenicillin media and harvested at an OD of 2. Serial dilutions of the initial transformation were plated to ensure sufficient library diversity (>50x coverage of 60,000 guides). Plasmid library was purified using NEB miniprep kit and pooled library was analyzed by Sanger sequencing.

Barcodes were then added to the gRNA plasmid library, targeting an average of four barcodes per guide RNA (240,000 barcodes in total). First, barcode DNA fragments were generated by annealing oligos RM504 and RM505 and extending the duplex using Q5 polymerase. The following thermocycler conditions were used to avoid amplification of any specific barcode sequence. Initial denaturation at 98 °C for 45s, six cycles of annealing and extension at 68 °C for 15s and 72 °C for 5s respectively, and finally a 12 °C hold. The barcode fragments were digested with BamHI and MfeI in order to eliminate barcode fragments that contain these restriction sites and then purified using a DNA Clean and Concentrator-5. The library of gRNA-expressing plasmids was then digested with AflII and the barcode DNA fragments were introduced by Gibson assembly. The assembly reaction was purified using a DNA Clean and Concentrator-5 then transformed into NEB^®^ 10-beta Competent E. coli (High Efficiency) according to manufacturer’s protocol. Transformation dilutions were plated to estimate total number of transformants, and in parallel inoculated at several dilutions directly into LB-carbenicillin media. Liquid culture corresponding to roughly 240,000 transformants was harvested at an OD of 2 and plasmid library was extracted using NEB miniprep kit. Reagents were scaled to accommodate one spin column for every 5mL of culture. This barcoded gRNA library was paired-end sequenced (see “Barcode to gRNA assignment” section) to assign barcodes with guide RNAs.

*P(HIS4)*-yECitrine was amplified from pNTI741 using primers RM501 and RM502, *P(PGK1)*-yECitrine was amplified from pNTI725 using primers RM501 and RM503, and two versions of estradiol-inducible P(*Z*)-mCherry with unique nucleotide identifiers were amplified with either RM522 and RM524 or RM523 and RM524. The barcoded gRNA plasmid library was digested with BamHI and the four PCR products were introduced into the digested library by Gibson assembly to create the four CiBER-seq libraries. These libraries were then transformed into ElectroMAX DH10B cells by electroporation.

Electroporated cells were inoculated directly into 500mL of LB-carbenicillin media and harvested at an OD of 2. Serial dilutions of the initial transformation were plated to ensure sufficient library diversity (>50x coverage of 60,000 guides). Plasmid library was purified using NEB miniprep kit and pooled library was analyzed by Sanger sequencing.

### Construction of yeast strains

All yeast strains were constructed by transforming PCR products or plasmids linearized by restriction enzyme digestion into yeast by standard lithium acetate transformation *(57)*. Selected, clonal transformants were confirmed by two genotyping PCRs, using primer pairs that amplify across the junction on either side of the integration. Genotyping reactions were performed with OneTaq Hotstart Polymerase (NEB #M0481S).

Yeast strains used in this study are listed in Table 3. Strains with *gcn2Δ* or *gcn4Δ* genotype were constructed by first amplifying the hygromycin resistance cassette from pNTI729 by PCR, with flanking sequence homologous to the 5′ and 3′ ends of the *GCN2* or *GCN4* coding sequence, using primers RM579 through RM582 for *gcn2Δ* and primers RM583 through RM586 for *gcn4Δ*. Primers RM589 and RM590 were used to genotype the upstream junction for *gcn2Δ* while RM591 and RM592 were used for the downstream junction. Primers RM593 and RM590 were used to genotype the upstream junction for *gcn4Δ* while RM591 and RM594 were used for the downstream junction.

Guide RNA expression cassettes were integrated using vectors from the EasyClone 2.0 toolkit for yeast genomic integration (55). Plasmids using backbone pCfB2189 were integrated by digesting with NotI, transforming this digestion, and selecting transformants on SCD -Leu plates. Plasmids using backbone pCfB2337 were integrated by digesting with NotI, transforming this digestion, plating transformations on non-selective YEPD media overnight, and then replica plating on hygromycin plates. Histidine prototrophy was restored by digesting pCfB2226 with NotI, transforming into the appropriate strain, and selecting transformants on SCD -His plates. Plasmid pNTI647, expressing dCas9 and tetR, was integrated as described in *(53)*. Integration of guide RNA expression cassettes was validated by amplifying across the upstream junction with RM527 and RM373 and across the downstream junction with RM374 and RM528. Integration of synthetic transcription factor variants were genotyped by amplifying across the upstream junction with RM372 and RM373 and across the downstream junction with RM374 and RM375.

### Media

LB media was prepared by dissolving LB medium capsules (MP Biomedicals #3002-31) in ultra-pure water and sterilized by autoclaving, according to manufacturer instructions. LB Media was supplemented with 50 $g/ml carbenicillin (Sigma #C1389) for antibiotic selection. YEPD was prepared by dissolving yeast extract (RPI # Y20026) and peptone (RPI #20241) in ultra-pure water, sterilizing by autoclaving, then supplementing with 2% final concentration of sterilized dextrose (Fisher Chemical #D16-500). Synthetic complete dropout (SCD) media was prepared by dissolving Yeast Nitrogen Base (BD Difco #291940), dextrose to 2% final concentration, and the appropriate drop-out mix in ultra-pure water, then sterilizing by filtration. SCD -Leu drop-out media was prepared using Drop-out Mix Synthetic Minus Leucine (US Biological #D9525), SCD -Ura drop-out media was prepared using Drop-out Mix Synthetic Minus Uracil (US Biological #D9535).

### qRT-PCR

RNA was isolated from yeast by acid phenol extraction *(58)*. Reverse transcription was carried out using Protoscript II reverse transcriptase (NEB #M0368L) with oligo-dT priming according to manufacturer instructions. Quantitative RT-PCR was carried out using DyNAmo qPCR mastermix (ThermoFisher

#F410L) according to manufacturer protocols, using a CFX96 Touch Real-Time PCR Detection System. Primers, provided in Table 5, were designed using Primer Blast and validated by using a cDNA dilution standard curve.

### Fluorescence measurements

Expression of yECitrine, under the control of the ZEM transcription factor, (Fig. 1) was monitored using a 96-well plate reader (Tecan SPARK^®^ Multimode Microplate Reader). Overnight cultures were back-diluted in 96-well round bottom plates to OD 0.05 in SCD -Ura selective media containing beta-estradiol and / or anhydrotetracycline as indicated. Fluorescence (excitation at 516nm and emission at 540nm) and OD600 was measured in triplicate every 15 minutes.

### Turbidostat continuous culture

Yeast populations transformed with plasmid libraries were inoculated into a custom turbidostat *(59)* and maintained in SCD -Ura media at a density corresponding to OD_600_ roughly 0.8. When growth rate reached a steady state (corresponding to a doubling time of roughly 90 minutes), a 50 ml pre-induction sample was collected. Guide RNA expression was then induced by injecting concentrated anyhydrotetracycline into both the growth chamber and turbidostat media reservoir to obtain a final concentration of 250 ng / ml of anyhydrotetracycline. Six doublings later (9 hours) a 50 mL post-induction sample was collected. For 3AT treatment, samples were induced by injecting concentrated 3AT into both the growth chamber and turbidostat media reservoir to obtain a final concentration of 90 mM 3AT. Two hours later, a post-induction sample was collected. For CiBER-seq experiments involving the ZEM transcription factor, media was supplemented with 8 nM beta-estradiol throughout the course of the experiment. Collected samples were pelleted by centrifugation at 4,000 × *g* for 5 minutes, the media was aspirated, and the pellets were stored at −80°C.

### Barcode sequencing

All PCR reactions were performed using Q5 polymerase (NEB M0491S) according to manufacturer protocols. PCR cycle numbers were adjusted as needed (between 6-12 cycles) to obtain adequate concentration of product for sequencing, while avoiding overamplification. DNA was purified using DNA Clean & Concentrator (Zymo #D4013) according to the manufacturer instructions. When necessary, AMPure XP beads (Beckman Coulter #A63881) were used to purify full-length DNA product. Size distributions and concentrations of the sequencing libraries were measured before sequencing using an Agilent TapeStation 2200 and High Sensitivity D1000 ScreenTape. To ensure proper clustering on the Illumina HiSeq4000, CiBER-seq libraries were either pooled with RNA-seq libraries and standard 2% PhiX, or individually with 10% PhiX. PCR products were pooled and sequenced on HISeq 4000 SR50 (Vincent J Coates Sequencing facility). Sequencing data accession numbers are available in Table 7.

### DNA library preparation

Plasmid DNA was extracted from yeast pellets using Zymo yeast miniprep kit (Zymo #D2004) according to manufacturer protocol, except as described below. Reagent volumes were scaled to accommodate roughly 250 million cells, as yeast pellets included 25 OD_600_ ml of yeast. Zymolyase concentration was increased to 1μL for every 10 million cells and the digestion time was doubled to ensure complete cell wall digestion.

Extracted barcode expression plasmid libraries derived from pNTI726 and pNTI728 backbones (Fig. 1) were prepared by PCR amplification of barcode sequences with primers that incorporated flanking TruSeq adapters. Barcode sequences were first amplified by PCR with RM411 and an i5 primer from NEBNext^®^ Multiplex Oligos (NEB #E7600S) for 12 cycles. PCR products were purified and concentrated using a DNA Clean and Concentrator-5, and then used as the template for an additional 6 cycles of PCR amplification with NEBNext i5 and i7 primers (NEB #E7600S).

To achieve a more linear amplification, all other CiBER-seq DNA libraries were prepared using an initial linear amplification by in vitro transcription. Extracted plasmid was linearized by restriction digestion with MfeI for *P(PGK1)* and *P(HIS4)* plasmid libraries (Fig. 2 through 4) or PvuII for *P(Z*) libraries (Fig. 5 and 6). The digestion product was purified using a DNA Clean and Concentrator-5, and used as the template for an IVT reaction using T7 HiScribe (NEB #E2040S) according to the manufacturer protocol for short transcripts, with an overnight incubation at 37°C. The reaction was then treated with DNase I to remove template DNA and RNA product was purified using an RNA Clean and Concentrator-5 (Zymo #R1016). Purified RNA was used as the template for reverse transcription using ProtoScript II according to manufacturer protocol, using sequence-specific primer RM511 for *P(PGK1)* and *P(HIS4)* plasmid libraries (Fig. 2 through 4) or RM546 for *P(Z)* libraries (Fig. 5 and 6). Reverse transcription cDNA product was treated for 30 mins with 0.5 μL RNAse A (ThermoFisher #EN0531) and 0.5 μL RNAse H (NEB #M0297S) to remove RNA, and then DNA was purified using a DNA Clean and Concentrator-5 column. The sequencespecific reverse transcription primers RM511 and RM546 incorporate the i7 priming site, and the i5 priming site is included in the plasmid. The final library was generated using 8 cycles of PCR amplification using NEBNext i5 and i7 dual index primers, and purified cDNA product as template. The resulting product was purified using a DNA Clean and Concentrator-5, and submitted for Illumina high throughput sequencing.

### RNA library preparation

Total RNA was harvested from yeast pellets as described above for qPCR analysis. Extracted RNA from libraries using the pNTI726 or pNTI728 backbones (Fig. 1) was used as the template for reverse transcription by ProtoScript II using sequence-specific primer RM411, according to the manufacturer instructions. Reverse transcription product was purified using a DNA Clean and Concentrator-5 and used as the template for PCR amplification using i5 and i7 dual indexing primers (NEB #E7600S).

In CiBER-seq libraries, the barcode is present in the opposite orientation. Reverse transcription was carried out using ProtoScript II according to the manufacturer protocol for oligo-dT priming. Reverse transcription product was treated with 0.5 μL RNAse A and 0.5 μL RNAse H for 30 min to remove RNA, and the DNA product was purified using a DNA Clean and Concentrator-5. The resulting cDNA product was first amplified by PCR for 6 cycles using primers RM511 and RM512 for *P(HIS4)* and *P(PGK1)* libraries, or primers RM512 and RM546 for *P(Z)* libraries. The PCR products were then purified and subsequently amplified in 8 cycles of PCR using NEBNext i5 and i7 dual indexing primers. PCR products were purified using a DNA Clean and Concentrator-5, and submitted for Ilumina high-throughput sequencing.

### Barcode to gRNA assignment

The sequencing library for paired-end barcode-to-gRNA assignment sequencing was constructed by PCR amplification from the barcoded gRNA library using primers RM506 and RM509 for 6 cycles using Q5 polymerase according to the manufacturer instructions. The resulting PCR product was purified using a DNA Clean and Concentrator-5 and then used as a template for a second, 15-cycle PCR using NEBNext i5 and i7 dual indexing primers. The PCR product was purified using a DNA Clean and Concentrator-5 and analyzed using an Agilent TapeStation 2200 and High Sensitivity D1000 ScreenTape. The PCR product library was then sequenced using 150+150 base paired-end sequencing on an Illumina MiSeq. Barcode to gRNA assignment sequencing data is available under accession SRR10327353.

### Single gRNA induction

Yeast strains expressing individual gRNAs were grown overnight in YEPD, then diluted to OD_600_ of 0.01 in -Leu media containing either 250 ng/ ml anhydrotetracycline, or no inducer. Yeast were harvested 12 hours later (after roughly 6 doublings), pelleted at 4,000 × *g* for 5 minutes, and pellets were frozen and stored at −80 °C for subsequent RNA extraction or protein isolation.

### Western blotting

Total protein was isolated from frozen yeast pellets by resuspending in 1.5 ml of 5% trichloroacetic acid and incubated at 4°C for <10 minutes. Protein was pelleted by centrifugation at 16,000 × *g* for 2 minutes and washed in 0.5 ml acetone, vortexed briefly, and collected by centrifugation for 5 min at 16,000 × *g*. Protein pellets were washed once more in 1. ml acetone, vortexed, and collected by centrifugation for 5 min at 16,000 × *g*. Pellets were dried overnight and then resuspended in 100 $l of freshly prepared protein breakage buffer (50 mM Tris•HCl pH 7.5, 1 mM EDTA, 3 mM DTT) containing 100$L of glass beads (BioSpec 11079105). Samples were vortexed in 5 cycles of one minute vortexing followed by one minute on ice. Lysates were transferred to new tubes and collected by centrifugation at 16,000 × g, then resuspended in 150 $l resuspension buffer (100 mM Tris•HCl pH 11, 3% SDS). Samples were boiled for 5 minutes then allowed to cool, and pelleted at 16,000 × g for 30 seconds. 120 $l of lysate was transferred to a new tube, and a BCA assay was performed to determine protein abundance. Equal amounts of total protein were loaded on a 4-12% polyacrylamide Bis-Tris gel (Thermo Scientific #NW04120BOX) and separated by electrophoresis in MOPS buffer at 200 V in a Bolt gel tank (Thermo Scientific #A25977) according to manufacturer instructions. Protein was then transferred to a nitrocellulose membrane (Thermo Scientific #88018) according to manufacturer guidelines. Membranes were blocked for 1 hour in TBST with 5% milk, washed with TBST three times for 10 minutes of shaking, and incubated with primary antibodies in TBST plus 0.5% milk overnight. Hexokinase loading control was probed with rabbit anti-Hxk2 (Rockland #100-4159) and phosphorylated eIF2α was detected using Phospho-eIF2a (Ser52) Polyclonal Antibody (Invitrogen #44-728G). Membranes were then washed and probed with secondary antibody anti-rabbit IgG, HRP-linked Antibody (Cell Signaling Technology #7074S). Membrane was imaged by Pierce ECL Western Blotting Substrate (ThermoScientific #32209) and the chemiluminescence measured on a FluorChem R (ProteinSimple).

### Sequencing data analysis

Sequencing data was processed using Cutadapt to remove constant adapter sequences and demultiplex libraries based on embedded nucleotide identifiers. The adapter sequences for each experimental dataset are provided in Table 6. The underlined sequence represents the library-specific nucleotide identifier.

For dual-guide experiments (Fig. 4), demultiplexing was not required as each pool derived from a single *P(HIS4)* library. For this reason, adapter trimming was performed by instead taking the first 25 nucleotides using fastx_trimmer.

Trimmed barcodes were then counted using the custom “bc-seqs” program, which collapses barcode variants separated by single-nucleotide mismatches (53). Using the “bc-tabulate” script, these barcode counts were then collected into a matrix, tabulating samples within an experiment. The *P(HIS4)* and *P(PGK1)* libraries were sequenced in two Illumina HiSeq4000 runs and the barcode counts for the runs were summed.

Barcodes were first filtered to remove those that lacked at least 32 counts in the pre-induction DNA sample for at least one replicate. The remaining barcodes were evaluated by differential activity analysis using a massively-parallel reporter assay linear model (mpralm)*(60)*. Barcode expression was compared between samples before and after gRNA induction, or before and after 3AT treatment. Analysis was performed using the aggregate = “sum” parameter to sum barcodes that corresponded to the same guide RNA and model_type = “indep_groups”, which treats the replicates as independent experiments. Output tables were merged with gene information from the Saccharomyces Genome Database. Scatterplots comparing log fold change values of individual gRNAs showing were first thresholded for guides in which *q* < 0.05 in at least one of the two expression analyses. Barcodes corresponding to no-guide plasmids were used as negative controls to determine the distribution of *p* values. Genes involved in amino acid biosynthesis, amino-acyl tRNA charging, ISR-controlled translation initiation, RNA polymerase III transcription, tRNA processing, or actin cytoskeletal arrangement were hand-curated and used to annotate volcano and scatter plots.

## Notes

#### Summary of Updates

Added reference to contemporaneous study employing similar approach: B. D. Alford, G. Valiant, O. Brandman, Genome-wide, time-sensitive interrogation of the heat shock response under diverse stressors via ReporterSeq. bioRxiv, doi.org:10.1101/2020.03.29.014845 (2020).

https://ingolia-lab.github.io/CiBER_seq

