## Supplemental Figures and Tables for "CiBER-seq dissects genetic networks by quantitative CRISPRi profiling of expression phenotypes"

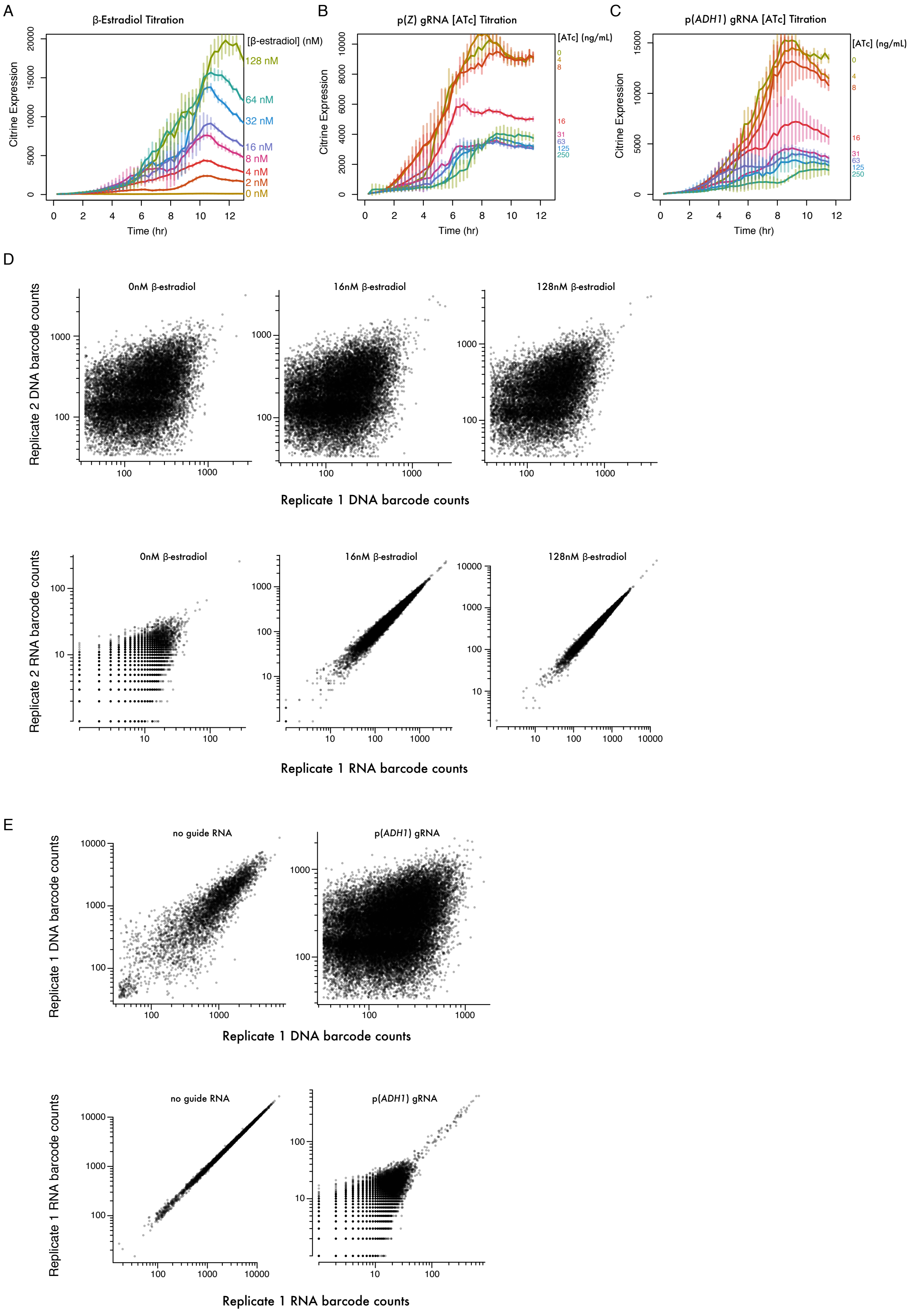
A
β-Estradiol Titration
[β-estradiol] (nM)
128 nM
64 nM
32 nM
16 nM
8 nM
4 nM
2 nM
0 nM
Citrine Expression
Time (hr)
B
p(Z) gRNA [ATc] Titration
[ATc] (ng/mL)
0
4
8
16
31
63
125
250
Citrine Expression
Time (hr)
C
p(ADH1) gRNA [ATc] Titration
[ATc] (ng/mL)
0
4
8
16
31
63
125
250
Citrine Expression
Time (hr)
D
0nM β-estradiol
16nM β-estradiol
128nM β-estradiol
Replicate 2 DNA barcode counts
Replicate 1 DNA barcode counts
0nM β-estradiol
16nM β-estradiol
128nM β-estradiol
Replicate 2 RNA barcode counts
Replicate 1 RNA barcode counts
E
no guide RNA
p(ADH1) gRNA
Replicate 1 DNA barcode counts
Replicate 1 DNA barcode counts
no guide RNA
p(ADH1) gRNA
Replicate 1 RNA barcode counts
Replicate 1 RNA barcode counts

**Fig. S1. Barcoded expression reporters accurately link guide RNAs with transcriptional phenotypes.**
(**A**) Fluorescence timecourse across estradiol titration demonstrates tunable activity of *P(Z)*.
(**B** and **C**) Fluorescence timecourse across anhydrotetracycline induction of guide RNAs targeting (B) *P(Z)* and (C) *P(ADH1)-ZEM* demonstrate tunable CRISPR interference with direct or indirect effects on reporter expression.
(**D** and **E**) Comparison between barcode sequence counts in replicate DNA and RNA sequencing libraries. Note that DNA libraries were constructed by PCR with no linear IVT amplification.

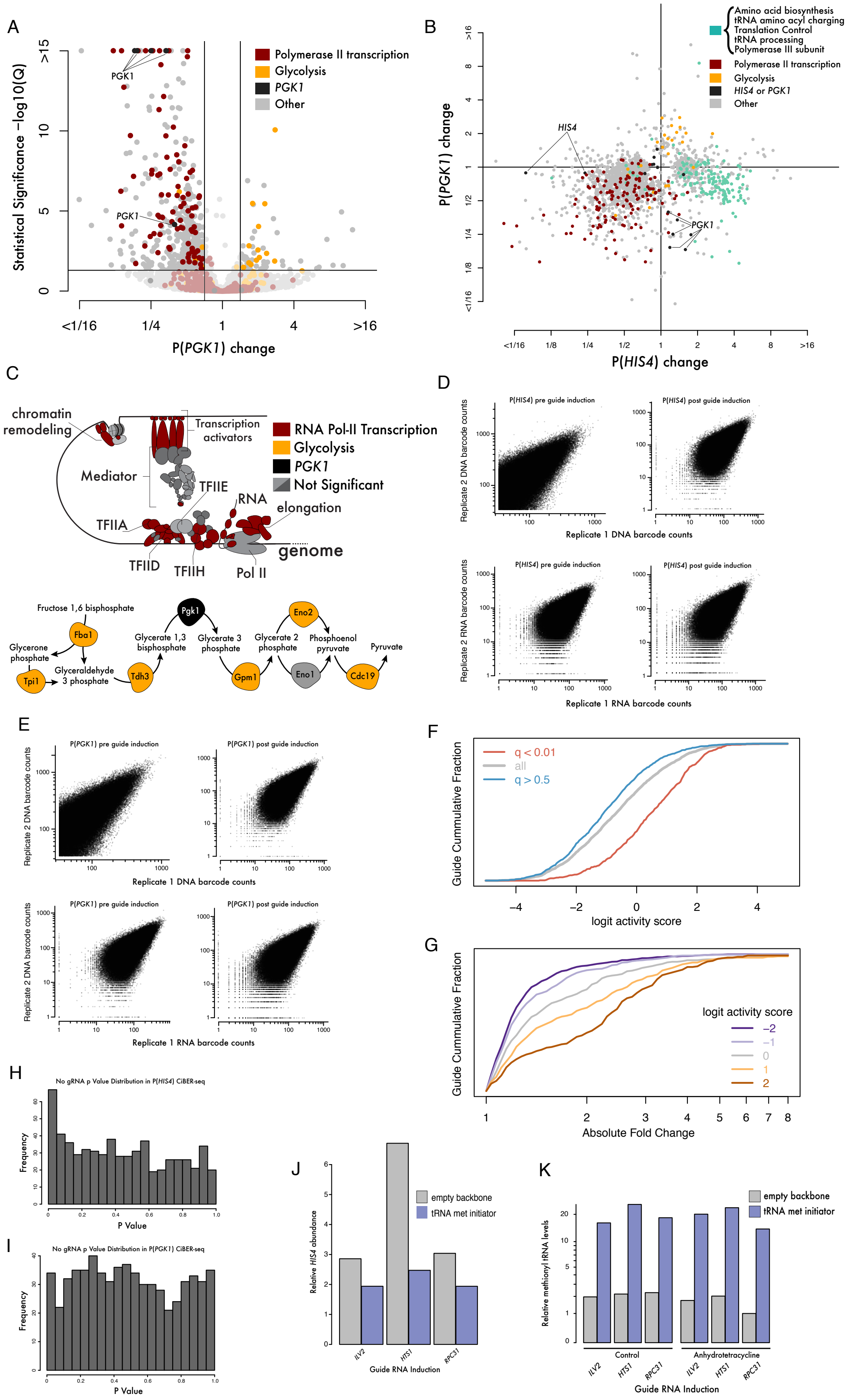
A
Statistical Significance −log10(Q)
PGK1
Polymerase II transcription
Glycolysis
PGK1
Other
P(PGK1) change
B
P(PGK1) change
P(HIS4) change
HIS4
PGK1
Amino acid biosynthesis
tRNA amino acyl charging
Translation Control
tRNA processing
Polymerase III subunit
Polymerase II transcription
Glycolysis
HIS4 or PGK1
Other
C
chromatin remodeling
Transcription activators
Mediator
TFIIE
RNA elongation
TFIIA
TFIID
TFIIH
Pol II
genome
RNA Pol-II Transcription
Glycolysis
PGK1
Not Significant
Fructose 1,6 bisphosphate
Fba1
Glycerone phosphate
Tpi1
Glyceraldehyde 3 phosphate
Tdh3
Glycerate 1,3 bisphosphate
Pgk1
Glycerate 3 phosphate
Gpm1
Glycerate 2 phosphate
Eno2
Eno1
Phosphoenol pyruvate
Cdc19
Pyruvate
D
P(HIS4) pre guide induction
P(HIS4) post guide induction
Replicate 2 DNA barcode counts
Replicate 1 DNA barcode counts
Replicate 2 RNA barcode counts
Replicate 1 RNA barcode counts
E
P(PGK1) pre guide induction
P(PGK1) post guide induction
Replicate 2 DNA barcode counts
Replicate 1 DNA barcode counts
Replicate 1 RNA barcode counts
F
q < 0.01
all
q > 0.5
Guide Cummulative Fraction
logit activity score
G
Guide Cummulative Fraction
logit activity score
Absolute Fold Change
H
No gRNA p Value Distribution in P(HIS4) CiBER-seq
Frequency
P Value
I
No gRNA p Value Distribution in P(PGK1) CiBER-seq
Frequency
P Value
J
Relative HIS4 abundance
empty backbone
tRNA met initiator
ILV2
HTS1
RPC31
Guide RNA Induction
K
Relative methionyl tRNA levels
empty backbone
tRNA met initiator
Control
Anhydrotetracycline
ILV2
HTS1
RPC31
Guide RNA Induction

**Fig. S2. CiBER-seq recapitulates known genetic regulators of integrated stress response and identifies new regulators related to tRNA insufficiency.**

(**A**) Genome-wide CiBER-seq analysis of housekeeping P(*PGK1*) promoter. Each point represents a different guide, colored according to the function of the target gene.

(**B**) Comparison of CiBER-seq profiles of P(*HIS4*) and P(*PGK1*) transcription, as in Fig. 2A, but colored as in (A).

(**C**) Functional analysis of guides with significant effects on *P(PGK1)* expression. Many guides that decrease *P(PGK1)* activity map to general RNA polymerase II transcription factors. Guides that increase *P(PGK1)* activity map to glycolysis.

(**D** and **E**) Comparison between barcode sequence counts in replicate DNA and RNA sequencing libraries for (D) *P(HIS4)* and (E) *P(PGK1)* barcodes.

(**F** and **G**) Among all guides targeting genes with ISR activation phenotypes, (F) those with $q < 0.01$ have larger logit guide activity scores than those with $q > 0.5$, and (G) those with a greater logit activity scores have larger effects on reporter expression.

(**H** and **I**) Histogram of unadjusted $p$ values for non-targeting barcodes in (H) *P(HIS4)* and (I) *P(PGK1)* CiBER-seq experiments. Uniform $p$-value distributions supports the null hypothesis used for significance testing of active guides.

(**J**) Analysis of *HIS4* mRNA levels by qPCR to assess ISR activation by CRISPRi knockdown of *ILV2*, *HTS1*, or *RPC31* in conjunction with initiator methionyl-tRNA overexpression. Initiator tRNA overexpression shows the same, partial rescue of *HIS4* transcriptional activation for each guide.

(**K**) Analysis of Met-tRNA levels verifies initiator methionyl-tRNA overexpression. Of note, *RPC31* knockdown decreases Met-tRNA abundance.

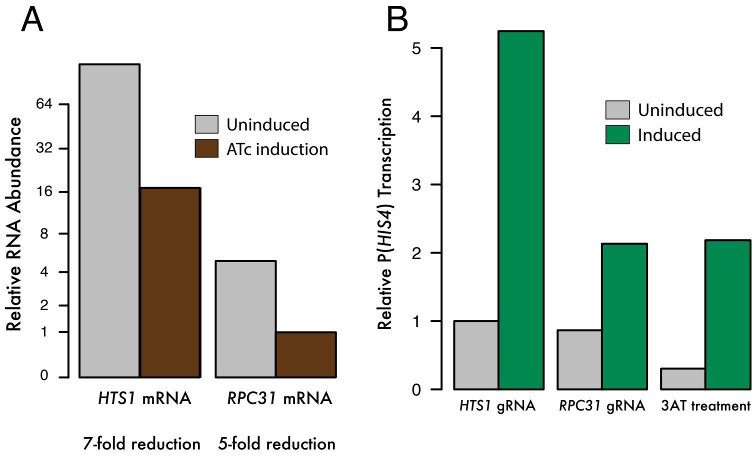


**Fig. S3. tRNA insufficiency activates *HIS4* transcription through a pathway that bypasses GCN2 kinase and eIF2α phosphorylation.**

(**A**) Endogenous *HTS1* and *RPC31* mRNA abundance, normalized to *ACT1* mRNA levels, demonstrates strong, inducible CRISPRi knockdown.

(**B**) Levels of *P(HIS4)* reporter transcript confirm the magnitude of ISR activation by *HTS1* knockdown, *RPC31* knockdown, or 3AT treatment.

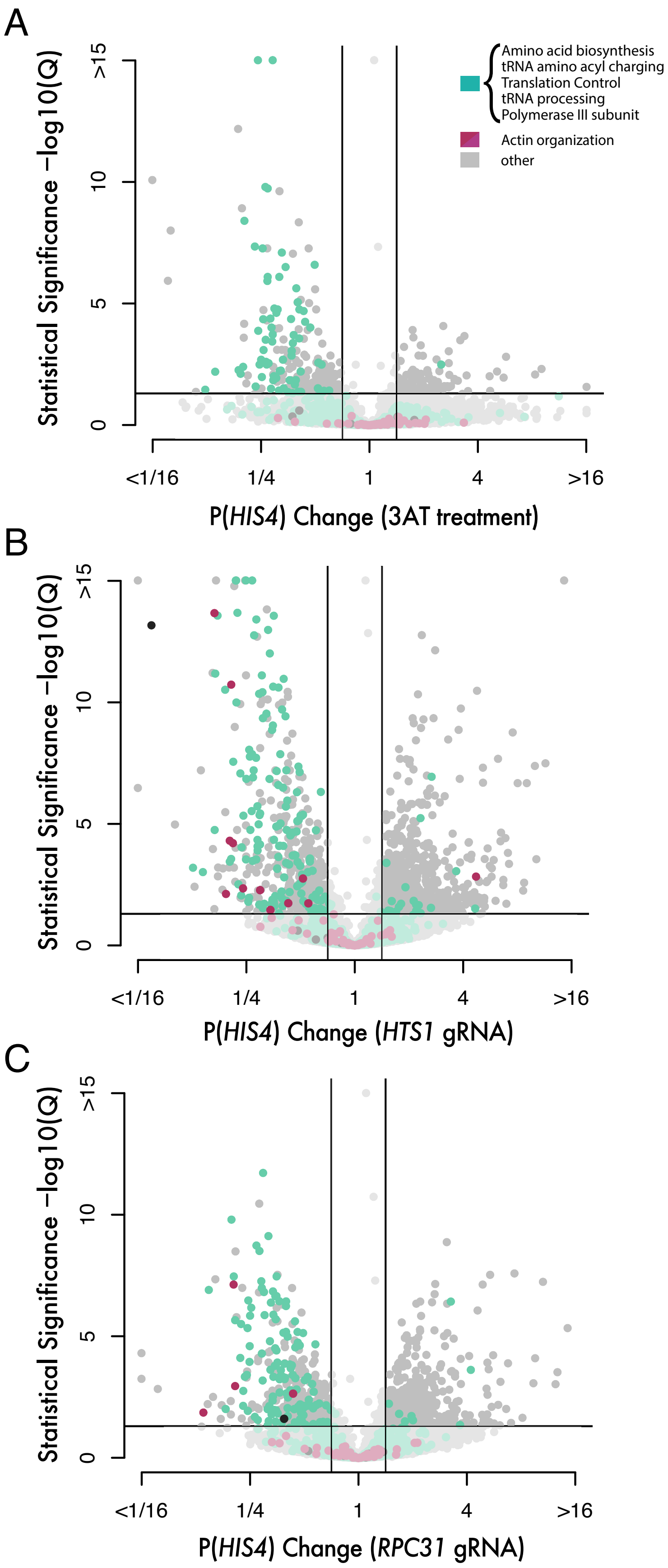
A
Amino acid biosynthesis
tRNA amino acyl charging
Translation Control
tRNA processing
Polymerase III subunit
Actin organization
other
Statistical Significance −log10(Q)
P(HIS4) Change (3AT treatment)
B
P(HIS4) Change (HTS1 gRNA)
C
P(HIS4) Change (RPC31 gRNA)

**Fig. S4. Perturbations of the *ARP2/3* complex prevent ISR activation by *HTS1* or *RCP31* knockdown.**
(**A** through **C**) Genome-wide P(*HIS4*) CiBER-seq profiles, showing (A) before and after 3AT treatment, (B) comparison of P(*HIS4*) expression after induction of one library guide versus after that same guide in conjunction with *HTS1* knockdown, and (C) same as (**B**), but in conjunction with *RPC31* knockdown. Each point represents a guide colored according to the function of the target gene.

A

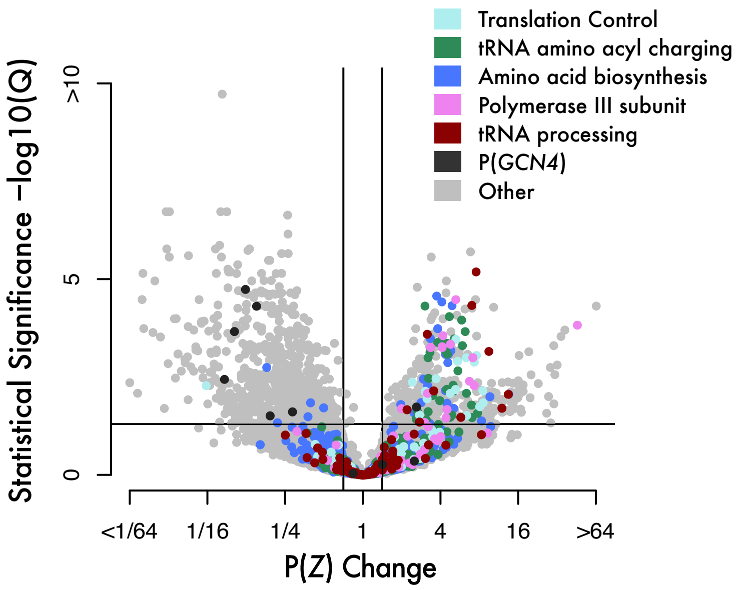


**Fig. S5. The GCN4 5′ leader sequence is an intrinsic biosensor of translation stress.**
(**A**) Indirect CiBER-seq profiling to measure gRNA effects on *GCN4* 5′ leader translation regulation as in Fig. 2A.

Table 1. Plasmids used in this study

| Plasmid | Description | Reference | Appearance in this work |
|---|---|---|---|
| pNTI601 | AddGene #73796 | *(19)* | |
| pNTI646 | gRNA expression in -ura CEN/ARS | This work | |
| pNTI660 | gRNA expression in -ura CEN/ARS | This work | |
| pNTI189 | (pKT0139 AddGene #8731) | *(53)* | |
| pNTI725 | p(*PGK1*) + yECitrine CGA in pNTI660 | This work | |
| pNTI642 | (pHES836 Addgene #89195) | *(17)* | |
| pNTI726 | Estradiol-inducible p(*Z*) + yECitrine CGA in pNTI660 | This work | Fig. 1A-1C |
| pNTI727 | Estradiol-inducible p(*Z*) + yECitrine CGA in pNTI660 with p(GAL1) targeting gRNA | This work | Fig. 1D, 1E |
| pNTI728 | Estradiol-inducible p(*Z*) + yECitrine CGA in pNTI660 with p(*ADH1*) targeting gRNA | This work | Fig. 1D, 1E, 1F |
| pNTI638 | (pHES795 Addgene #87943) | *(17)* | |
| pCfB2337 | HygR easyclone 2.0 (Addgene #67555) | *(54)* | |
| pCFB2226 | spHis5 easyclone 2.0 (Addgene #67533) | *(54)* | |
| pNTI729 | p(*ADH1*)-ZEM Transcription Factor in pCfB2337 | This work | Fig. 1A-1F |
| pNTI730 | p(*PGK1*)-ZEM Transcription Factor in pCfB2337 | This work | |
| pCfB2189 | Leu2 easyclone 2.0 (Addgene #67532) | *(54)* | |
| pNTI731 | *ILV2* gRNA in pCfB2189 | This work | Fig. 3C-3E |

| | | | |
|---|---|---|---|
| pNTI732 | *HTS1* gRNA in pCfB2189 | This work | Fig. 3C-3E, 4A, 4C |
| pNTI733 | *RPC31* gRNA in pCfB2189 | This work | Fig. 3C-3E, 4B, 4C |
| pNTI734 | *NUP133* gRNA in pCfB2189 | This work | Fig. 5C |
| pNTI735 | *NUP145* gRNA in pCfB2189 | This work | Fig. 5C |
| pNTI736 | *ULP1* gRNA in pCfB2189 | This work | Fig. 5C |
| pNTI737 | *UBC9* gRNA in pCfB2189 | This work | Fig. 5C |
| pRS426 | 2-micron plasmid (ATCC #77107) | *(55)* | fig. S2J, S2K |
| pNTI738 | met initiator tRNA overexpression in pRS426 | this work | fig. S2J, S2K |
| pNTI739 | p(*PGK1*) + *GCN4*(CDS) + ZEM(TF) in pCfB2337 | This work | Fig. 5A-5C |
| pNTI740 | p(*GCN4*) + *GCN4*(UTR) + ZEM(TF) in pCfB2337 | This work | Fig. 6A, 6B |
| pNTI741 | P(*HIS4*) + yECitrine template in pNTI660 | This work | |
| pNTI742 | Modular parent vector | This work | Fig. 2-6 |

Table 2. Primers used in this study

| Primer | Primer sequence |
|---|---|
| NM636 | ggctgggaacgaaactctgggagctgcgattggca |
| NM637 | gccttattttaacttgctatttctagctctaaaac. |
| NM717 | ccctacacgttcgctatgcaaaccctggcgttaccc |
| NM724 | aaaccctggcgttacccaacttaatcgccttgcagca |
| NI553 | CGGATCCTGTAGCCCTAGACTTGATAGCCATGACTTCAACTCAAGACGCACAG |
| NI554 | TTCTTCACCTTTAGACATGTTAATTAATTTGTTGTAAAAAGTAGATAATTACTTCCTTG |
| RM151 | TTTTACAACAAATTAATTAAATGTCTAAAGGTGAAGAATTATTCACTGGTGTTGTC |
| RM152 | TTATTTGTACAATTCATCCATACCATGGGTAATAC |
| RM317 | TTCAAGCATACCAAGTTGGTgatctgccaattgaacataacatggtag |
| RM318 | ctaccatgttatgttcaattggcagatcACCAACTTGGTATGCTTGAA |
| RM319 | ttttgctcgggggatctgccTGAGCTCGAATTGATCCTGTAGCC |
| RM320 | GGCTACAGGATCAATTCGAGCTCAggcagatcccccgagcaaaa |
| RM321 | ctgggagctgcgattggcagTCGCGAgttttagagctagaaatagc |
| RM323 | GCGCCCCTCCACGGGATATCGGCTAAACACTCTTTCCCTACACGACGCTCTTCCGATCTG |
| RM324 | CTTATTTAGAAGTGGCGCGCTCTGTTGATAACTCCGGATCCAGATCGGAAGAGCGTCGTG |
| RM348 | gatctgccTGAGCTCGACCGGTttatattgaattttcaaaaattcttactttttttttgg |
| RM349 | TCTTCACCTTTAGACATTTGGATCCtatagttttttctccttgacgttaaagtatagagg |

| | |
|---|---|
| RM352 | gaagatcgcgtcagctgaagGGCAACCAAACCCATACATCGG |
| RM353 | aaccagcaccgtcacctgctgcggcgactgtggcagg |
| RM354 | tttccctgccacagtcgccgcagcaggtgacggtgctgg |
| RM355 | cgacctgcagcgtacgaaggatccacatcaaaaggcctctaggttcc |
| RM372 | CGGGTGGCCAAAGCAAAGG |
| RM373 | ctcaggtatagcatgaggtcgctc |
| RM374 | gatccactagtggcctatgcaccc |
| RM375 | GAAGCCAAAAGACCAGAGTAGAGGCC |
| RM383 | ctgggagctgcgattggcagagggaacgagaacaatgacg |
| RM384 | gctatttctagctctaaaaccgtcattgttctcgttccct |
| RM389 | ctgggagctgcgattggcagtccttgacgttaaagtatagg |
| RM390 | gctatttctagctctaaaacctatactttaacgtcaaggactgcc |
| RM391 | ACGCTCTTCCGATCTNNNNNNNNNNACTGCNNNNNNNNNNGGAGTTATCAACAGAGCGCG |
| RM392 | ACGCTCTTCCGATCTNNNNNNNNNNCTGAGNNNNNNNNNNGGAGTTATCAACAGAGCGCG |
| RM393 | ACGCTCTTCCGATCTNNNNNNNNNNGACTANNNNNNNNNNGGAGTTATCAACAGAGCGCG |
| RM394 | ACGCTCTTCCGATCTNNNNNNNNNNTGACTNNNNNNNNNNGGAGTTATCAACAGAGCGCG |
| RM396 | CGCGCTCTGTTGATAACTCC |
| RM399 | ACGCTCTTCCGATCTNNNNNNNNNNAAGCTNNNNNNNNNNGGAGTTATCAACAGAGCGCG |

| | |
|---|---|
| RM411 | GTGACTGGAGTTCAGACGTGTGCTCTTCCGATCT CGCGCTCTGTTGATAACTCC |
| RM417 | acgctcgaagatcgcgtcagctgaagGACTTCAACTCAAGACGCACAG |
| RM418 | cgggtacccattttattgctagccATTTGTTGTAAAAAGTAGATAATTACTTCCTTGATG |
| RM459 | ctgggagctgcgattggcagCCTAGgttttagag |
| RM460 | gctatttctagctctaaaacCTAGGctgccaatc |
| RM489 | ctgccTGAGCTCTAATACGACTCACTATAGCAGATCTATATTACCCTGTTATCCCTAGCG |
| RM490 | AAGATCGGAAGAGCGTCGTGTAGGGAAAGAGTGTAAGCTTGCGCGCCACTTCTAAATAAG |
| RM491 | aactaccatgttatgttcaattggcagatcGGATCCCTTAAGATCGGAAGAGCGTCGTG |
| RM499 | tcgggggatctgccTGAGCTCGACCGGTTAGGCAGTCGAACTGACTCTAATAG |
| RM500 | ACCTTTAGACATTTGCTAGCTATTCAGAAAAAAAATTTTGTAAACTATTGTATTAC |
| RM501 | GAGATCCAGTCACTCGGaactgTTACTATTTGTACAATTCATCCATACCATGG |
| RM502 | tatgttcaattggcagatcGTAGGCAGTCGAACTGACTCTAATAGTG |
| RM503 | tatgttcaattggcagatcGGACTTCAACTCAAGACGCACAGATATTATAAC |
| RM504 | CGACGCTCTTCCGATCTNNNNNNNNNNNNNNNNNNNNNNNNNGAGATCCAGTCACTCGGG |
| RM505 | gttcaattggcagatcGGATCCCGAGTGACTGGATCTC |
| RM506 | GCTTATTTAGAAGTGGCGCGCAAG |
| RM509 | TTCAGACGTGTGCTCTTCCGATCTagccttattttaacttgctatttctagctctaaaac |
| RM511 | GTGACTGGAGTTCAGACGTGTGCTCTTCCGATCTTGGTCCAGTCTTGTTACCAGACAACC |

| | |
|---|---|
| RM512 | CGCGCAAGCTTACACTCTTTCCC |
| RM513 | cgctcgaagatcgcgtcagctgaagTACATCTTGAAAAAAAAAGATGAAAAATTTCC |
| RM514 | tacccattttattgctagccTTTATTTGTATTTAATTTATTTTCTTGAGCAGACAAATTG |
| RM515 | CTACTTTTTACAACAAATggctagcATGTCCGAATATCAGCCAAGTTTATTTG |
| RM516 | aagcatatgggcgggtacccatAGATCCAGAACCGCGTTCGCCAACTAATTTCTTTAATC |
| RM517 | gaagatcgcgtcagctgaag AAGACGTTATCGGGGCGCGCCTGGG |
| RM518 | gaagatcgcgtcagctgaag GTATACCATGACTGATTACGTACAT |
| RM519 | tcgacctgcagcgtacgaagcccatttttagcatgtaaatataaagagaaacc |
| RM522 | GAGATCCAGTCACTCGGccttgCTATTTGTACAATTCATCCATACCACCAGTAG |
| RM523 | GAGATCCAGTCACTCGGtcagaCTATTTGTACAATTCATCCATACCACCAGTAG |
| RM524 | tatgttcaattggcagatcGttatattgaattttcaaaaattcttactttttttttgg |
| RM527 | TTGAACATTTTATGATTTCTTGCTCTTGG |
| RM528 | CAAGAACAATAATAAGGTTTCCTGAAGG |
| RM546 | GTGACTGGAGTTCAGACGTGTGCTCTTCCGATCTTCAAGTTGGACATCACCTCCCAC |
| RM579 | AATAATTTTCCGTTCCCCTTAACACATACTATGTATAAggcggcagaagaagtaacaaag |
| RM580 | AACTGATGCGTTATAGCGCCGCACAGATCTTTAAAGGCttgggtgcataggccactagtg |
| RM581 | TAAGATTTTATAAGCATTGATTTTTTTTTCAATAATTTTCCGTTCCCCTTAACACATAC |
| RM582 | TAAATATAGTATCTAAGACATTGTATATACTTTACCTTTAACTGATGCGTTATAGCGCCG |
| RM583 | ATTTGTCTGCTCAAGAAAATAAATTAAATACAAATAAAggcggcagaagaagtaacaaag |
| RM584 | GAATGAAATAAAAAATATAAAATAAAAGGTAAATGAAAttgggtgcataggccactagtg |

| | |
|---|---|
| RM585 | TTACTAAAGTTTTGTTTACCAATTTGTCTGCTCAAGAAAATAAATTAAATACAAATAAAg |
| RM586 | CTATTTCGTTATACACGAGAATGAAATAAAAAATATAAAATAAAAGGTAAATGAAAttgg |
| RM589 | GATGTAGGTAGTTTCTTAGGAAGCAGTTG |
| RM590 | aatgattatacatggggatgtatggg |
| RM591 | agttgttctattttaatcaaatgttagcgtg |
| RM592 | GGAGCTAGAGGAGCCTCACACAAC |
| RM593 | GAAAGAGAAAATTTATTTTCCCTTATTAATTAAAGTCC |
| RM594 | AATAAATAATGCTCGCGTGGCG |
| RM636 | tatcgataagcttgatatcgAATATCCATGGGAGAAATCACCATC |
| RM637 | tggatcccccgggctgcaggCAAGGGTCAGTCAATCACTTAACCAC |
| RM639 | gaagatcgcgtcagctgaag GCTCTTTTATAAAGTCCCTATGCCC |
| RM641 | gaagatcgcgtcagctgaag TCACTAGGGGTGTCGGGTCTCCACC |
| RM642 | gaagatcgcgtcagctgaag GTTGTGACGCAAGTGCCGTAGGTCC |
| RM643 | gaagatcgcgtcagctgaag GCGTTAGGTAGAGGCGGGCAAACGT |
| RM644 | gaagatcgcgtcagctgaag CGCCTACGCGGCGGCACCATGCGAG |

Table 3. Yeast strains used in this study

| Strain | Genotype | Reference |
|---|---|---|
| BY4741 | MATa his3Δ1 leu2Δ LYS2 met15Δ ura3Δ0 | ATCC #201388 |
| NIY416 | MATa his3Δ1 leu2Δ LYS2 met15Δ ura3Δ0 [dCas9 tetR, G418R] | McGlincy et al |
| NIY443 | MATa his3Δ1 leu2Δ LYS2 met15Δ ura3Δ0 [dCas9 tetR, G418R] [p(ADH1)-ZEM Transcription Factor, HygR] | This work |
| NIY444 | MATa his3Δ1 leu2Δ LYS2 met15Δ ura3Δ0 [dCas9 tetR, G418R] [spHis5] | This work |
| NIY445 | MATa his3Δ1 leu2Δ LYS2 met15Δ ura3Δ0 [dCas9 tetR, G418R] [p(PGK1)-(GCN4 CDS-ZEM TF fusion), HygR] [spHis5] | This work |
| NIY446 | MATa his3Δ1 leu2Δ LYS2 met15Δ ura3Δ0 [dCas9 tetR, G418R] [p(GCN4)-(GCN4 5'UTR-ZEM TF), HygR] [spHis5] | This work |
| NIY447 | MATa his3Δ1 leu2Δ LYS2 met15Δ ura3Δ0 [dCas9 tetR, G418R] [spHis5] [ILV2 gRNA, LEU2] | This work |
| NIY448 | MATa his3Δ1 leu2Δ LYS2 met15Δ ura3Δ0 [dCas9 tetR, G418R] [spHis5] [ILV2 gRNA, LEU2] [gcn2Δ HygR] | This work |
| NIY449 | MATa his3Δ1 leu2Δ LYS2 met15Δ ura3Δ0 [dCas9 tetR, G418R] [spHis5] [ILV2 gRNA, LEU2] [gcn4Δ HygR] | This work |
| NIY450 | MATa his3Δ1 leu2Δ LYS2 met15Δ ura3Δ0 [dCas9 tetR, G418R] [spHis5] [HTS1 gRNA, LEU2] | This work |
| NIY451 | MATa his3Δ1 leu2Δ LYS2 met15Δ ura3Δ0 [dCas9 tetR, G418R] [spHis5] [HTS1 gRNA, LEU2] [gcn2Δ HygR] | This work |
| NIY452 | MATa his3Δ1 leu2Δ LYS2 met15Δ ura3Δ0 [dCas9 tetR, G418R] [spHis5] [HTS1 gRNA, LEU2] [gcn4Δ HygR] | This work |
| NIY453 | MATa his3Δ1 leu2Δ LYS2 met15Δ ura3Δ0 [dCas9 tetR, G418R] [spHis5] [RPC31 gRNA, LEU2] | This work |

| | | |
|---|---|---|
| NIY454 | MATa his3Δ1 leu2Δ LYS2 met15Δ ura3Δ0 [dCas9 tetR, G418R] [spHis5] [RPC31 gRNA, LEU2] [gcn2Δ HygR] | This work |
| NIY455 | MATa his3Δ1 leu2Δ LYS2 met15Δ ura3Δ0 [dCas9 tetR, G418R] [spHis5] [RPC31 gRNA, LEU2] [gcn4Δ HygR] | This work |
| NIY456 | MATa his3Δ1 leu2Δ LYS2 met15Δ ura3Δ0 [dCas9 tetR, G418R] [p(PGK1)-(GCN4 CDS-ZEM TF fusion), HygR] [spHis5] [NUP133 gRNA, LEU2] | This work |
| NIY457 | MATa his3Δ1 leu2Δ LYS2 met15Δ ura3Δ0 [dCas9 tetR, G418R] [p(PGK1)-(GCN4 CDS-ZEM TF fusion), HygR] [spHis5] [NUP145 gRNA, LEU2] | This work |
| NIY458 | MATa his3Δ1 leu2Δ LYS2 met15Δ ura3Δ0 [dCas9 tetR, G418R] [p(PGK1)-(GCN4 CDS-ZEM TF fusion), HygR] [spHis5] [ULP1 gRNA, LEU2] | This work |
| NIY459 | MATa his3Δ1 leu2Δ LYS2 met15Δ ura3Δ0 [dCas9 tetR, G418R] [p(PGK1)-(GCN4 CDS-ZEM TF fusion), HygR] [spHis5] [UBC9 gRNA, LEU2] | This work |

Table 4. Targeting guide RNA sequences (underlined is gRNA scaffold, bold is 20nt guide)

| Gene target promoter | **Guide Sequence** + gRNA scaffold | Reference |
|---|---|---|
| *GAL1* | **tccttgacgttaaagtatag**gttttagagctagaaatagcaagttaaaataaggctagtccgttatcaacttgaaaaagtggcaccgagtcggtgc | This work |
| *ADH1* | **agggaacgagaacaatgacg**gttttagagctagaaatagcaagttaaaataaggctagtccgttatcaacttgaaaaagtggcaccgagtcggtgc | This work |
| *ILV2* | **GAACTTGTATTTCTCTTATC**gttttagagctagaaatagcaagttaaaataaggctagtccgttatcaacttgaaaaagtggcaccgagtcggtgc | This work |
| *HTS1* | **CTATTCTAAAGTAACACATT**gttttagagctagaaatagcaagttaaaataaggctagtccgttatcaacttgaaaaagtggcaccgagtcggtgc | This work |
| *RPC31* | **TTCTATAGGTTGCTGCGATG**gttttagagctagaaatagcaagttaaaataaggctagtccgttatcaacttgaaaaagtggcaccgagtcggtgc | This work |
| *NUP133* | **TTCCTAAATGGTGTATGTTT**gttttagagctagaaatagcaagttaaaataaggctagtccgttatcaacttgaaaaagtggcaccgagtcggtgc | This work |
| *NUP145* | **AACAACAATCACATCACCAT**gttttagagctagaaatagcaagttaaaataaggctagtccgttatcaacttgaaaaagtggcaccgagtcggtgc | This work |
| *ULP1* | **TGAATTTTGAAAATAAAAAG**gttttagagctagaaatagcaagttaaaataaggctagtccgttatcaacttgaaaaagtggcaccgagtcggtgc | This work |
| *UBC9* | **TAAATGTTTGTTATGGAGAC**gttttagagctagaaatagcaagttaaaataaggctagtccgttatcaacttgaaaaagtggcaccgagtcggtgc | This work |

Table 5. qPCR primers used in this study

| Primer | Primer sequence | Gene target/primer purpose |
|---|---|---|
| RM529 | ACCGCTAACAATACCTGGGC | *URA3*_forward |
| RM530 | ATGGAGGGCACAGTTAAGCC | *URA3*_reverse |
| RM531 | GAAATGCAAACCGCTGCTCA | *ACT1*_forward |
| RM532 | TACCGGCAGATTCCAAACCC | *ACT1*_reverse |
| RM533 | GGATGCTGGCATTGAAGCTG | *HTS1*_forward |
| RM534 | GCCGCACTAACCAATTCACC | *HTS1*_reverse |
| RM535 | TTCCATCCATCCCATTGCCC | *RPC31* _forward |
| RM536 | TTTGCGTTTACCGCTCTTGC | *RPC31*_reverse |
| RM595 | AAGGAGCTTTCTTGGGAGGC | *HIS4*_forward |
| RM596 | GCTCAAAGCCTTCTGCACAC | *HIS4*_reverse |
| RM679 | TAGCGCCGCTCGGTTTCGAT | initiator-met-tRNA_first strand cDNA |
| RM680 | GTGGCGCAGTGGAAG | initiator-met-tRNA_forward |
| RM681 | CGAGGACATCAGGGTTATG | initiator-met-tRNA_reverse |
| RM341 | ACACTCTTTCCCTACACGACGC | TruSeq barcode_forward |
| RM342 | TCTGTTGATAACTCCGGATCCAG | TruSeq barcode_reverse |
| RM537 | CATGGTCTTCTTCTGCATTACG | mCherry_forward |
| RM538 | GACTACTTGAAGCTGTCCTTC | mCherry_reverse |

Table 6. Adapter sequences

| Library | Trimmed 5' Adapter Sequence | Trimmed 3' Adapter Sequence |
|---|---|---|
| Paired-end barcode gRNA assignment (barcode) | | GAGATCCAGTCACTCGGGATCCgatc tgccaattgaacataacatggtagt |
| Paired-end barcode gRNA assignment (gRNA) | aagttaaaataaggct | |
| p(*Z*)-yEcitrine | | GGAGTTATCAACAGAGCGCGAGAT CG |
| P(*HIS4*)-yEcitrine | | GAGATCCAGTCACTCGGaactgTTAC |
| P(*PGK1*)-yEcitrine | | GAGATCCAGTCACTCGGtcgatTTAC |
| P(*Z*)-mCherry for GCN4(CDS) screen | | GAGATCCAGTCACTCGGCCTTGCTA T |
| P(*Z*)-mCherry for GCN4(UTR) screen | | GAGATCCAGTCACTCGGTCAGACT AT |

Table 7. SRA accession numbers and related information for each sample used in this study

| **Accession** | Title | BioSample | Submission | SRA.filename |
|---|---|---|---|---|
| **SRR10357613** | CRISPRi minimal media growth, day 0, left culture | SAMN13150799 | SUB6475866 | 11-20_R2_PCR_B_S11_L001_R1_001.fastq.gz |
| **SRR10357612** | CRISPRi minimal media growth, day 0, right culture | SAMN13150799 | SUB6475866 | 11-20_R2_PCR_C_S12_L001_R1_001.fastq.gz |
| **SRR10357609** | CRISPRi minimal media growth, day 2, left culture | SAMN13150799 | SUB6475866 | exp_bcode_dup1_S5_L001_R1_001.fastq.gz |
| **SRR10357608** | CRISPRi minimal media growth, day 2, right culture | SAMN13150799 | SUB6475866 | exp_bcode_dup2_S6_L001_R1_001.fastq.gz |
| **SRR10352291** | CiBER-Seq P(HIS4) with RPC31 RNA Pre-induction replicate 1 | SAMN13134381 | SUB6472309 | RNA_RPC31_Pre_S7_R1_001.fastq.gz |
| **SRR10352293** | CiBER-Seq P(HIS4) with HTS1 RNA Post-induction replicate 1 | SAMN13134380 | SUB6472309 | RNA_HTS1_Post_S6_R1_001.fastq.gz |
| **SRR10352294** | CiBER-Seq P(HIS4) with HTS1 DNA Post-induction replicate 1 | SAMN13134380 | SUB6472309 | IVT_HTS1_Post_S2_R1_001.fastq.gz |
| **SRR10352292** | CiBER-Seq P(HIS4) with RPC31 DNA Pre-induction replicate 1 | SAMN13134381 | SUB6472309 | IVT_RPC31_Pre_S3_R1_001.fastq.gz |
| **SRR10352302** | CiBER-Seq P(HIS4) with HTS1 DNA Pre-induction replicate 1 | SAMN13134379 | SUB6472309 | IVT_HTS1_Pre_S1_R1_001.fastq.gz |
| **SRR10352290** | CiBER-Seq P(HIS4) with RPC31 DNA Post-induction replicate 1 | SAMN13134382 | SUB6472309 | IVT_RPC31_Post_S4_R1_001.fastq.gz |
| **SRR10352301** | CiBER-Seq P(HIS4) with HTS1 RNA Pre-induction replicate 1 | SAMN13134379 | SUB6472309 | RNA_HTS1_Pre_S5_R1_001.fastq.gz |
| **SRR10352289** | CiBER-Seq P(HIS4) with RPC31 RNA Post-induction replicate 1 | SAMN13134382 | SUB6472309 | RNA_RPC31_Post_S8_R1_001.fastq.gz |
| **SRR10352298** | CiBER-Seq P(HIS4) with RPC31 DNA Pre-induction replicate 2 | SAMN13134385 | SUB6472309 | RPC31_DNA_pre.fastq.gz |

| | | | | |
|---|---|---|---|---|
| **SRR10352299** | CiBER-Seq P(HIS4) with HTS1 RNA Post-induction replicate 2 | SAMN13134384 | SUB6472309 | HTS1_RNA_post.fastq.gz |
| **SRR10352300** | CiBER-Seq P(HIS4) with HTS1 DNA Post-induction replicate 2 | SAMN13134384 | SUB6472309 | HTS1_DNA_post.fastq.gz |
| **SRR10352297** | CiBER-Seq P(HIS4) with RPC31 RNA Pre-induction replicate 2 | SAMN13134385 | SUB6472309 | RPC31_RNA_pre.fastq.gz |
| **SRR10352295** | CiBER-Seq P(HIS4) with RPC31 RNA Post-induction replicate 2 | SAMN13134386 | SUB6472309 | RPC31_RNA_post.fastq.gz |
| **SRR10352288** | CiBER-Seq P(HIS4) with HTS1 DNA Pre-induction replicate 2 | SAMN13134383 | SUB6472309 | HTS1_DNA_pre.fastq.gz |
| **SRR10352296** | CiBER-Seq P(HIS4) with RPC31 DNA Post-induction replicate 2 | SAMN13134386 | SUB6472309 | RPC31_DNA_post.fastq.gz |
| **SRR10352287** | CiBER-Seq P(HIS4) with HTS1 RNA Pre-induction replicate 2 | SAMN13134383 | SUB6472309 | HTS1_RNA_pre.fastq.gz |
| **SRR10352193** | CiBER-Seq 5'UTR+CDS RNA Post-induction Left | SAMN13134388 | SUB6472302 | RNA_CU_PostL_S58_L007_R1_001.fastq.gz |
| **SRR10352191** | CiBER-Seq 5'UTR+CDS RNA Pre-induction Right | SAMN13134389 | SUB6472302 | RNA_CU_PreR_S65_L008_R1_001.fastq.gz |
| **SRR10352195** | CiBER-Seq 5'UTR+CDS RNA Pre-induction Left | SAMN13134387 | SUB6472302 | RNA_CU_PreL_S57_L007_R1_001.fastq.gz |
| **SRR10352189** | CiBER-Seq 5'UTR+CDS RNA Post-induction Right | SAMN13134390 | SUB6472302 | RNA_CU_PostR_S66_L008_R1_001.fastq.gz |
| **SRR10352190** | CiBER-Seq 5'UTR+CDS DNA Post-induction Right | SAMN13134390 | SUB6472302 | IVT2_CUpostR_S106_L003_R1_001.fastq.gz |
| **SRR10352194** | CiBER-Seq 5'UTR+CDS DNA Post-induction Left | SAMN13134388 | SUB6472302 | IVT2_CUpostL_S19_L008_R1_001.fastq.gz |
| **SRR10352192** | CiBER-Seq 5'UTR+CDS DNA Pre-induction Right | SAMN13134389 | SUB6472302 | IVT2_CUpreR_S105_L003_R1_001.fastq.gz |
| **SRR10352196** | CiBER-Seq 5'UTR+CDS DNA Pre-induction Left | SAMN13134387 | SUB6472302 | IVT2_CUpreL_S18_L008_R1_001.fastq.gz |
| **SRR10336437** | CiBER-Seq P(HIS4)+P(PGK1) DNA Post-induction Left | SAMN13088882 | SUB6454012 | IVT_PostL_S19_L008_R1_001.fastq.gz, IVT_PostL_S52_L008_R1_001.fastq.gz |

| | | | | |
|---|---|---|---|---|
| **SRR10336442** | CiBER-Seq P(HIS4)+P(PGK1) RNA Pre-induction Left | SAMN13088880 | SUB6454012 | RNA_PreL_S23_L008_R1_001.fastq.gz, RNA_PreL_S54_L008_R1_001.fastq.gz |
| **SRR10336434** | CiBER-Seq P(HIS4)+P(PGK1) RNA Post-induction Right | SAMN13088883 | SUB6454012 | RNA_PostR_S26_L008_R1_001.fastq.gz, RNA_PostR_S57_L008_R1_001.fastq.gz |
| **SRR10336436** | CiBER-Seq P(HIS4)+P(PGK1) RNA Post-induction Left | SAMN13088882 | SUB6454012 | RNA_PostL_S25_L008_R1_001.fastq.gz, RNA_PostL_S56_L008_R1_001.fastq.gz |
| **SRR10336439** | CiBER-Seq P(HIS4)+P(PGK1) DNA Pre-induction Right | SAMN13088881 | SUB6454012 | IVT_PreR_S18_L008_R1_001.fastq.gz, IVT_PreR_S51_L008_R1_001.fastq.gz |
| **SRR10336435** | CiBER-Seq P(HIS4)+P(PGK1) DNA Post-induction Right | SAMN13088883 | SUB6454012 | IVT_PostR_S20_L008_R1_001.fastq.gz, IVT_PostR_S53_L008_R1_001.fastq.gz |
| **SRR10336443** | CiBER-Seq P(HIS4)+P(PGK1) DNA Pre-induction Left | SAMN13088880 | SUB6454012 | IVT_PreL_S17_L008_R1_001.fastq.gz, IVT_PreL_S50_L008_R1_001.fastq.gz |
| **SRR10336438** | CiBER-Seq P(HIS4)+P(PGK1) RNA Pre-induction Right | SAMN13088881 | SUB6454012 | RNA_PreR_S24_L008_R1_001.fastq.gz, RNA_PreR_S55_L008_R1_001.fastq.gz |
| **SRR10336441** | CiBER-Seq P(HIS4)+P(PGK1) DNA Post-induction +3-AT Right | SAMN13088885 | SUB6454012 | IVT_3AT_R_S22_L008_R1_001.fastq.gz |
| **SRR10336433** | CiBER-Seq P(HIS4)+P(PGK1) DNA Post-induction +3-AT Left | SAMN13088884 | SUB6454012 | IVT_3AT_L_S21_L008_R1_001.fastq.gz |
| **SRR10336440** | CiBER-Seq P(HIS4)+P(PGK1) RNA Post-induction +3-AT Right | SAMN13088885 | SUB6454012 | RNA_3AT_R_S28_L008_R1_001.fastq.gz |
| **SRR10336432** | CiBER-Seq P(HIS4)+P(PGK1) RNA Post-induction +3-AT Left | SAMN13088884 | SUB6454012 | RNA_3AT_L_S27_L008_R1_001.fastq.gz |

| | | | | |
|---|---|---|---|---|
| **SRR10327353** | barcode-to-gRNA assignment | SAMN13088718 | SUB6453931 | mod_bc_gRNA_R1.fastq.gz, mod_bc_gRNA_R2.fastq.gz |
